# Robust temporal map of human *in vitro* myelopoiesis using single-cell genomics

**DOI:** 10.1101/2021.11.17.469005

**Authors:** Clara Alsinet, Maria Primo, Valentina Lorenzi, Andrew J Knights, Carmen Sancho-Serra, Jong-Eun Park, Beata S Wyspianska, David F Tough, Damiana Alvarez-Errico, Daniel J Gaffney, Roser Vento-Tormo

## Abstract

Myeloid cells have a central role in homeostasis and tissue defence. Characterising the current *in vitro* protocols of myelopoiesis is imperative for their use in research and immunotherapy as well as for understanding the early stages of myeloid differentiation in humans. Here, we profiled the transcriptome of more than 400k cells and generated a robust molecular map of the differentiation of human induced pluripotent stem cells (iPSC) into macrophages. By integrating our *in vitro* datasets with *in vivo* single-cell developmental atlases, we found that *in vitro* macrophage differentiation recapitulates features of *in vivo* yolk sac hematopoiesis, which happens prior to the appearance of definitive hematopoietic stem cells (HSC). During *in vitro* myelopoiesis, a wide range of myeloid cells are generated, including erythrocytes, mast cells and monocytes, suggesting that, during early human development, the HSC-independent immune wave gives rise to multiple myeloid cell lineages. We leveraged this model to characterize the transition of hemogenic endothelium into myeloid cells, uncovering poorly described myeloid progenitors and regulatory programs. Taking advantage of the variety of myeloid cells produced, we developed a new protocol to produce type 2 conventional dendritic cells (cDC2) *in vitro*. We found that the underlying regulatory networks coding for myeloid identity are conserved *in vivo* and *in vitro*. Using genetic engineering techniques, we validated the effects of key transcription factors important for cDC2 and macrophage identity and ontogeny. This roadmap of early myeloid differentiation will serve as an important resource for investigating the initial stages of hematopoiesis, which are largely unexplored in humans, and will open up new therapeutic opportunities.

---

Macrophages perform a variety of functions, ranging from tissue homeostasis to immune surveillance and from the response to infection to the resolution of inflammation^1–4^. They originate during both development and adulthood and acquire specific functions when they seed tissues^5^. To what extent cellular ontogeny and tissue microenvironment influence macrophage identity is poorly understood in humans. Despite the commonalities within mammals, there are important differences from rodent models^6^. Establishing and characterising the current human *in vitro* models is essential in order to fully exploit their potential in answering key biological questions and using them as novel therapeutic tools^7^.

During development, myeloid cells originate from at least two waves of progenitors: a first wave involving myeloid-biased progenitors from the yolk sac (YSMP) and a second wave through the definitive hematopoietic stem cells (HSC)^5,8^. YSMP are thought to appear during the first two weeks of development in humans and are responsible for producing primordial blood^5^. HSC are not generated until 3–4 post-conceptional weeks (PCW) in the gonad-aorta-mesonephros. HSC and myeloid progenitors derived from YSMP colonise the liver, making this fetal organ the main site of hematopoiesis until mid-pregnancy^9^. During the second trimester of development, HSC migrate to the bone marrow, the only remaining site of hematopoiesis in adulthood^10^. In mice, myeloid progenitors derived from the yolk-sac (YS) generate a wide range of myeloid cells, including monocytes and neutrophils, and are thought to be the main contributors to this lineage during fetal development^11^. In humans, we have limited knowledge about how the YS myeloid progenitors expand, proliferate and differentiate, and a poor understanding of the regulatory mechanisms involved.

*In vitro* models of macrophage differentiation hold promise to not only answer these biological questions but also to become therapeutic tools, particularly for immunotherapies. Macrophages derived from human induced pluripotent stem cells (iPSC) show tissue-resident phenotypes^12^ and are an attractive alternative to adult monocyte-derived macrophage cultures^13,14^. A current protocol, developed by van Wilgenburg et al., is a straightforward, feeder-free process done in 3 steps using between 1 and 3 cytokines at constant concentrations^15^. It provides long-term, scalable production of macrophage precursors without fluorescence-activated cell sorting (FACS), since the cells of interest continuously expand and detach from the culture^16^. Despite this being an established *in vitro* macrophage model, the exact intermediate populations that arise during the protocol are unclear^12^. This restricts its applications for iPSC manipulation (i.e., genetic screens) and limits our true understanding of the final cells produced. Thus, a thorough analysis of the cell identities and dynamics emerging during *in vitro* differentiation is imperative to fully exploit this technology.

Single-cell transcriptomics is a powerful tool for evaluating current *in vitro* models in relation to their *in vivo* counterparts^17,18^. Here, we profiled the single-cell transcriptome and open chromatin data of >400k and >70k cells, respectively, during iPSC–myeloid differentiation with the van Wilgenburg protocol^15^. We provide a roadmap of cell states emerging during iPSC–macrophage differentiation, along with their ontogeny and underlying transcription factor (TF) networks. We found that iPSC–macrophage differentiation accurately maps fetal myelopoiesis in the YS and generates a wide range of cell types, from endoderm to megakaryocytes and mast cells. We demonstrate the adaptability of the current *in vitro* protocol to produce alternative myeloid cells and induce distinct cell states by modifying the media used. Finally, we validate the effects of key TFs related to myeloid cell identity using the CRISPR-Cas9 genome editing of relevant genes involved in inflammatory diseases. Altogether, our study demonstrates that macrophage differentiation from iPSC is a robust system to study the early stages of myelopoiesis in humans, which would not be accessible otherwise, and produces macrophages able to polarise and acquire definitive tissue-resident identities.

## In vitro myelopoiesis has features of human yolk sac myelopoiesis

We profiled the full differentiation of iPSC into macrophages from 6 individuals using single-cell RNA sequencing (scRNAseq) and single-cell ATAC sequencing (scATACseq) **(Fig. 1A, Supplementary Table 1)**. The differentiation protocol consists of 3 steps: i) spin-embryoid body (EB) formation from day 1 to 4, ii) EB myeloid differentiation from day 5 onwards (the latest sample used in this study is from day 31), and iii) macrophage differentiation using non-adherent cells from day 31 (day 31 to day 31 + 7) **(Fig. 1A)**. To characterise the robustness of our results, we generated two independent scRNAseq datasets. In the first dataset (referred to as our Discovery dataset), we multiplexed scRNAseq data from 3 donors at 20 timepoints **(Fig. 1A-B)**. In the second dataset (hereafter, our Validation dataset), we multiplexed scRNAseq and scATACseq data from 6 donors at 7 and 6 time points, respectively **(Fig 1A-B)**. The three donors from the Discovery dataset were also used in the Validation dataset, thus generating biological replicates of those three lines.

**Fig 1.**
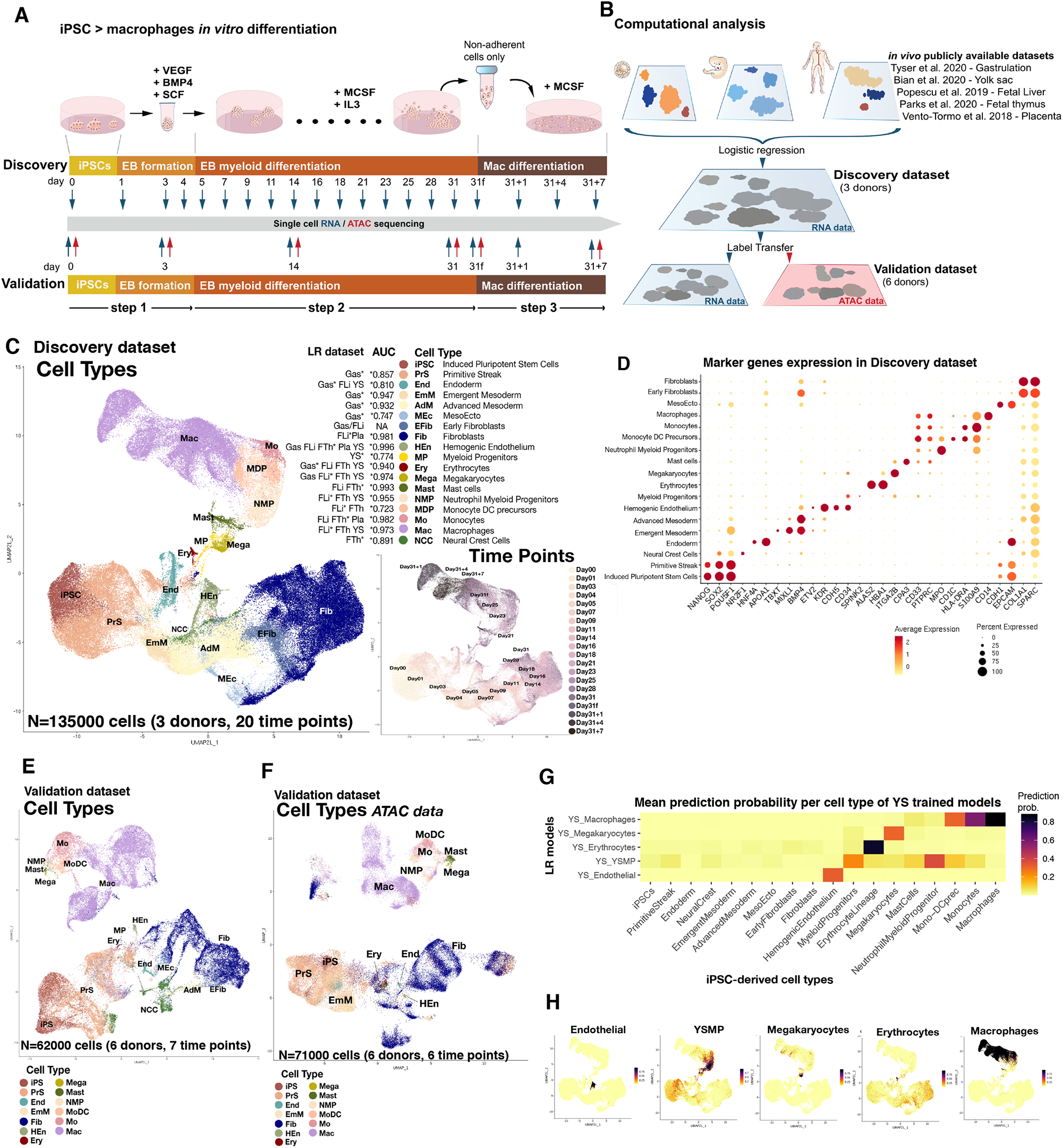
iPSC macrophage differentiation produces a range of fetal myeloid and stromal cells. **A**, Schematic illustration of the *in vitro* differentiation protocol from iPSC to macrophages highlighting the time points where samples were collected for scRNAseq and scATACseq profiling. The protocol was repeated twice to generate the Discovery and Validation datasets. **B**, Diagram summarising the computational workflow used for cell-type annotation of the single-cell datasets generated with the differentiation protocol. Briefly, LR models were used to annotate the Discovery scRNAseq dataset based on publicly available *in vivo* datasets of human gastrulation (Gas)^22^, yolk sac (YS)^6^, fetal liver (including skin and kidney) (FLi)^21^, fetal thymus (FTh)^20^ and placenta (Pla)^19^. Then, cell type annotations were transferred from the Discovery dataset to the scRNAseq and scATACseq Validation dataset. **C**, UMAP projections of the Discovery scRNAseq data (n = 135,000) from 3 cell lines and 20 time points labeled by cell type. For each cell type, we report the *in vivo* datasets supporting the annotation and the AUC on the Discovery dataset for the LR models trained on the *in vivo* dataset with a * mark. (right) UMAP projections of the Discovery dataset labeled by time point. **D**, Dot plot showing canonical markers for each of the cell types identified in the Discovery dataset. Colors depict the mean gene expression and dot size the percentage of cells expressing each marker. **E**, UMAP projections of the scRNAseq Validation dataset (n = 62,000) from 6 cell lines and 7 time points labeled by cell type. **F**, UMAP projections of the scATACseq Validation dataset (n = 71,000) from 6 cell lines and 6 timepoints labeled by cell type. **G**, Heatmap showing the mean logistic regression models’ predicted probabilities of the YS hematopoietic cell types^6^ for each of the cell types in the Discovery scRNAseq dataset. **H**, UMAP projections of the scRNAseq Discovery data coloured by the logistic regression models’ predicted probabilities of the YS hematopoietic cell types^6^. iPSC, induced pluripotent stem cell; EB, embryoid body; Mac, macrophage; LR, logistic regression; UMAP, uniform manifold approximation and projection; AUC, area under the curve; ATAC, assay for transposase-accessible chromatin; YS, yolk sac; YSMP, yolk sac myeloid-biased progenitors.

After quality control, the Discovery dataset contained a total of 135,000 cells **(Fig. 1C, Fig. S1)**. To annotate the cell types in an unbiased manner, we built logistic regression (LR) classifiers trained on publicly available single-cell transcriptomics datasets and projected the data into our iPSC–macrophage differentiation dataset **(Fig. 1B)**. We used multiple human embryonic (including gastrulation) and fetal datasets to train our classifiers^6,19–22^. Those datasets included the main fetal hematopoietic organs: yolk sac, liver and thymus (**Supplementary Table 2)**. Cell type labels were assigned based on the mean LR prediction probability of each cell cluster **(Fig. 1C, Fig. S2, Supplementary Table 3)**. Standard marker gene expression analysis further supported the cell type annotation obtained with LR **(Fig. 1D)**. Cell type label transfer^23^ from the Discovery dataset into the Validation dataset confirmed the presence of the same cellular subsets in both datasets **(Fig. 1E, Fig. S1, Supplementary Table 4)**. Most cell types were also recovered in the scATACseq dataset **(Fig. 1F, Supplementary Table 4**).

The majority of cells at the initial EB formation stage (the first 4 days) correspond to cell states present during gastrulation **(Fig. 1C)**. We found primitive streak-like cells, emergent and advanced mesoderm, and the initial appearance of hemogenic endothelium. Despite using cytokines that induce hematopoietic mesoderm (EB formation, **Fig. 1A**), we also observed two subsets of cells related to other germ layers (i.e., neural crest and endoderm, **Fig. 1C**).

During EB myeloid differentiation (5–31 days), the myeloid and stromal cell compartments emerged **(Fig. 1C)**. The myeloid populations produced *in vitro* include a wide range of cell types, such as erythrocytes, megakaryocytes, mast cells, neutrophil myeloid progenitors (NMP), monocyte-DC precursors (MDP), monocytes and macrophages **(Fig. 1C)**. The *in vivo* counterparts for these cells are found in the fetal liver and thymus. We did not find any cluster in our iPSC–macrophage dataset that corresponded to the HSC found in the human developing liver dataset **(Fig. S2B)**. Instead, there is a distinct cluster of myeloid progenitors (MP) that express CD34 and *SPINK2* but not the HOXA genes, which are required to generate definitive HSC^24,25^ **(Fig. 1D, Fig. S3A)**. The MP have a high prediction probability for the YSMP-trained model generated with the droplet-based embryonic YS dataset **(Fig. 1G-H, Fig. S2C)**, suggesting *in vitro* myelopoiesis recapitulates YS differentiation. Trained models with *in vivo* YSMP and macrophages captured more than one cell type within the *in vitro* dataset **(Fig. 1G-H)**. To explore this further, we did the opposite exercise: we trained models on our cell types defined *in vitro* and projected them onto the *in vivo* YS dataset. As expected, the *in vitro* MP LR model had a high probability for the *in vivo* YSMP cells **(Fig. S3B-C)**. In addition, we found a subset of cells within the original YSMP cluster that had a high prediction probability for the *in vitro* NMP-trained model. There was also a subset of *in vivo* macrophages that were identified by the MDP-trained model **(Fig. S3B-C)**. Using the LR results, we annotated the NMP and MDP cell types within the droplet-based embryonic YS dataset **(Fig. S3B)**.

To quantitatively characterise the fetal-like signature of the macrophages obtained with this protocol, we projected data from both adult and fetal macrophages into our dataset. We trained an LR classifier using macrophages from the human decidual–placental interface, a unique tissue setting that includes both adult/maternal monocyte-derived macrophages and fetal/placental YS-derived macrophages^19^, which allows us to overcome any potential issue with integrating fetal and adult datasets. YS-derived fetal macrophages (Hofbauer cells) had a higher mean prediction probability for iPSC-derived macrophages than for any of the adult macrophage subtypes identified in the placenta **(Fig. S3D-E)**. In line with this, LR models trained on fetal-like FOLR2+ and SPP1+ tumor-associated macrophages (TAM) from a hepatocellular carcinoma dataset^26^ show a high prediction probability for our iPSC-derived macrophages at distinct time points **(Fig. S3F-G)**. This indicates that macrophages produced in the iPSC protocol have a strong fetal phenotype, and this could be relevant for their application as *in vitro* TAM models.

Altogether, we show that *in vitro* iPSC differentiation to macrophages produces a plethora of myeloid cell types but lacks HSC, thus recapitulating yolk sac differentiation. Our map will be available at www.HiPImmuneatlas.org.

### Trajectory analysis and underlying regulatory programs

The generation of myeloid populations, including myeloid precursors, is controlled by several regulatory elements that shape transcriptional programs, including TFs, epigenetic regulators and post-transcriptional mechanisms^27^. We set out to reconstruct the main developmental pathways underlying *in vitro* hematopoiesis and the regulatory networks mediating them. The high-throughput single-cell approach used, along with the high density of time points collected and using scVelo^28^ for trajectory analysis, allowed us to reconstruct all the differentiation paths giving rise to the wide range of cell types observed **(Fig. 2A, Fig. S4A)**. In parallel, we set out to compare the underlying regulatory programs mediating such transitions *in vivo* and *in vitro*. To this end, we measured TF activities by looking at the expression of consensus TF targets^29^ **(Fig. 2B)**.

**Fig 2.**
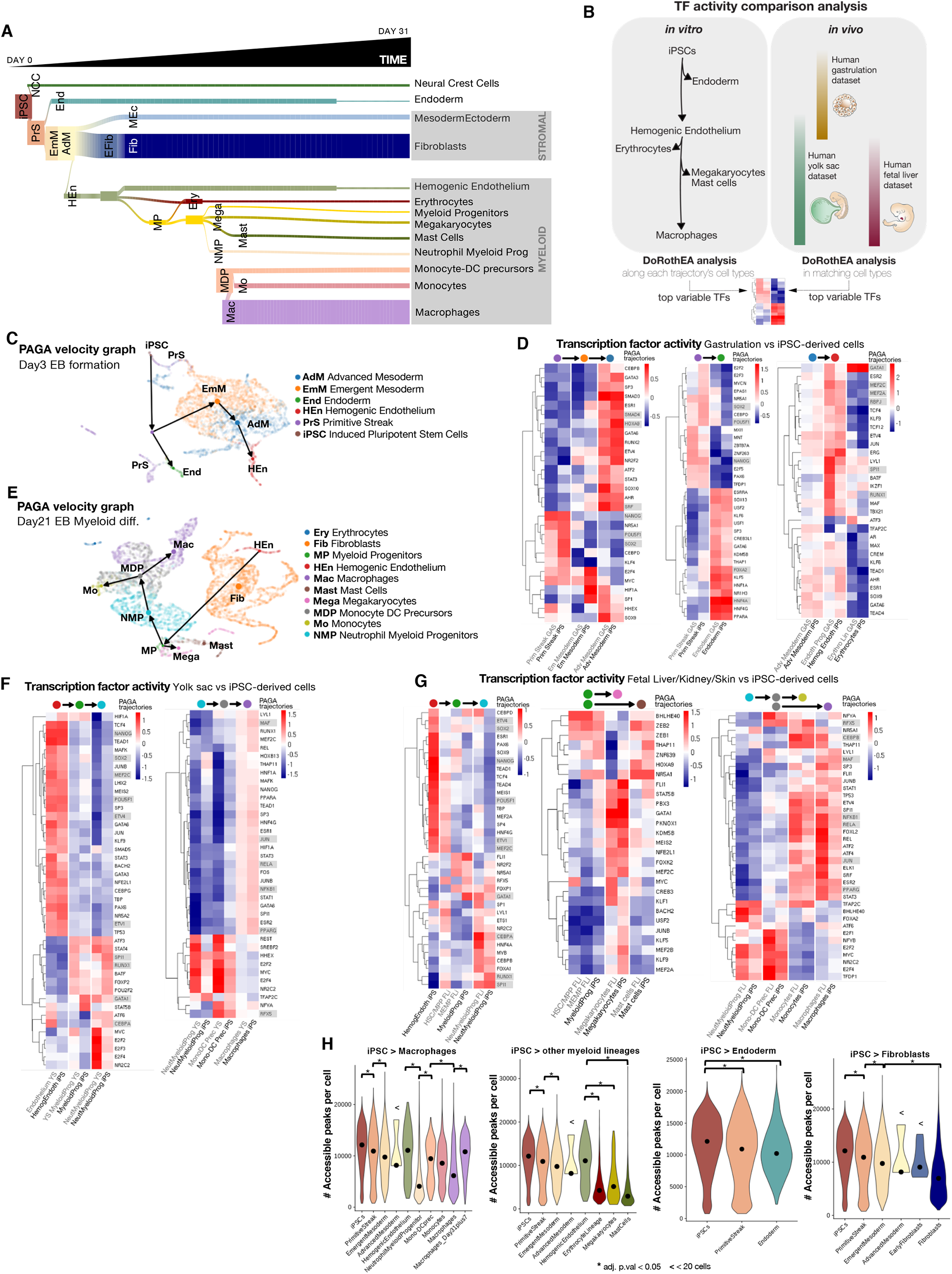
Cell population dynamics. **A**, Diagram illustrating the dynamic emergence of the different cell types over the course of the *in vitro* differentiation protocol (Discovery dataset). **B**, Schematic representation of the computational workflow used to compare transcription factor (TF) dynamics *in vivo* and *in vitro*. Briefly, TF activities were computed at branching points along the *in vitro* differentiation trajectory (Discovery dataset) and were compared to TF activities in matched cell types in the *in vivo* human yolk sac^6^, gastrulation^22^ and fetal liver^21^ datasets. **C**, RNA velocity and PAGA graph abstraction of the cells found at day 3 (Embryoid Body (EB) formation) of the differentiation protocol (Discovery dataset) showing the developmental relationships between the cell types. **D**, Transcription factor activities computed with DoRothEA for the identified cell types present at day 3 of the *in vitro* differentiation protocol and matched cell types in the *in vivo* gastrulation dataset^22^, relevant TF discussed in the text are highlighted in grey. **E**, RNA velocity and PAGA graph abstraction of the cells present at day 21 (EB myeloid differentiation) of the differentiation protocol (Discovery dataset) showing the developmental relationships between the cell types. **F**, Transcription factor activities computed with DoRothEA for the identified cell types present at day 21 of the *in vitro* differentiation protocol and matched cell types in the *in vivo* yolk sac dataset^6^, relevant TF discussed in the text are highlighted in grey. **G**, Same analysis as **F** with matched cell types in the *in vivo* fetal liver, skin and kidney dataset^21^. **H**, Violin plots showing the number of accessible peaks per cell type in the scATACseq Validation dataset. Each panel considers the cell types present in a distinct lineage (macrophages, other myeloid, endoderm, fibroblasts). iPSC, induced pluripotent stem cells; ATAC, assay for transposase-accessible chromatin.

In step 1 of *in vitro* differentiation, iPSC differentiate into the primitive streak, which subsequently gives rise to either endoderm or emergent and advanced mesoderm **(Fig. 2C)**. Later, advanced mesoderm can differentiate into hemogenic endothelium, which is the precursor of myeloid cells **(Fig. 2C)**. Primitive streak and mesoderm are transient populations that disappear by day 16, while endoderm and hemogenic endothelium remain stable until at least day 31 **(Fig. 2A, Fig. S4A)**. The transition from mesoderm to hemogenic endothelium has also been reported during the gastrulation period^30^. TF activity analysis shows high conservation of the TF modules in these earlier stages of development **(Fig. 2D, Supplementary Table 5)**. There is decreased activity of pluripotency TFs (POU5F1, NANOG and SOX2) when cells differentiate into mesoderm and endoderm. As expected, in both *in vivo* and *in vitro* settings, mesoderm activates SMAD3, HOXA9 or SRF while endoderm activates FOXA2 and HNF4A. Later, hemogenic endothelium activates TFs relevant for hematopoiesis including RUNX1, SPI1, RBPJ, MEF2A and MEF2C **(Fig. 2D)**. GATA1 is also activated in this transition but it shows the highest levels in erythrocytes **(Fig. 2D)**.

*In vitro*, myelopoiesis starts very early in the EB formation stage, but the wide range of myeloid cell types appear almost simultaneously starting at day 14, and they all endure at least until day 31, the latest time point of the EB myeloid differentiation phase collected **(Fig. 2A, Fig. S4A)**. At days 9–11, the first MP arise, followed by the appearance of erythrocytes on days 11–14, and full myelopoiesis is achieved on days 16–18. In addition to myeloid cells, advanced mesoderm also differentiates into an intermediate stage of early fibroblasts (day 7), giving rise to fibroblasts by day 9 **(Fig. S4A)**. Trajectory analysis on the sample at day 21 using scVelo reconstructs all myelopoiesis differentiation steps. Hemogenic endothelial cells, derived from the mesoderm, can differentiate into MP, which in turn, give rise to both megakaryocytes and NMP **(Fig. 2E, Fig. S4A)**. NMP give rise to MDP, which differentiate into either monocytes or macrophages **(Fig. 2E)**. The differentiation pathway of macrophages through MP, thus bypassing the monocytes, is consistent with the first waves of myelopoiesis emerging in the YS^5^.

Throughout all stages of myelopoiesis, we consistently found high similarity between the regulatory programs activated *in vivo* (embryonic YS and fetal liver) and *in vitro* (iPSC-derived cells) **(Fig. 2F-G, Supplementary Table 5)**. The transition from hemogenic endothelium to MP is characterized by the activation of TFs such as RUNX1, SPI1 and GATA1 **(Fig. 2F-G)**. The MP to NMP transition has further activation of SPI1 and CEBPA. On the contrary, a large number of endothelial TFs, such as SOX2 and the ETV family, pluripotency factors, such as POU5F1 and NANOG, or lymphoid lineage-promoting factors, such as MEF2C, are inactivated^31^. MEF2C is a TF characteristic of definitive HSC that drives lymphoid fate choice^31^, yet the *in vitro* MP have low MEF2C activity, which further supports the HSC-independent hematopoiesis profile of this system **(Fig. 2F)**.

Further differentiation towards monocytes and macrophages is also characterized by shared transcriptional programs between the iPSC-derived model and both its YS and fetal liver counterparts. Among the few TFs specifically activated in MDP from MP is RFX5, which regulates MHC-II transcription and is responsible for a rare hereditary immunodeficiency^32^. We also observed activation of TFs controlling inflammatory programs in monocytes and macrophages, such as JUN, RELA and NFKBI **(Fig 2F-G)**. Notably, we found key similarities in iPSC-derived macrophages compared to their *in vivo* counterparts in the YS and fetal liver datasets that account for the establishment of the myeloid identity through TFs such as MAF^33^ and CEBPB^34^ **(Fig 2F-G)**. There are also similarities that potentially underlie the basis for tissue-specific macrophage programs, such as the alveolar macrophage program represented by the activation of the tissue-specific TF PPARG^35,36^ **(Fig 2F-G)**.

Finally, we evaluated the chromatin accessibility dynamics of iPSC–macrophage differentiation. The number of accessible cell peaks decreased alongside the trajectories identified, suggesting a more restrictive chromatin landscape as cells differentiate **(Fig. 2H)**. An exception to this is the hemogenic endothelium. These cells have a median of 11074 accessible peaks per cell, which is higher than that of their mesoderm progenitors (‘Emergent Mesoderm’ n = 9759, ‘Advanced Mesoderm’ n = 8158), yet they share a similar number of expressed genes (‘Emergent Mesoderm’ n = 3463, ‘Advanced Mesoderm’ n = 2725, ‘Hemogenic endothelium’ n = 2514). Another exception is the macrophages after the macrophage differentiation phase (‘Macrophages_Day31plus7’), which have more accessibility peaks than do the macrophages collected from the EB myeloid differentiation phase (‘Macrophages’ n = 6142 vs ‘Macrophages_Day31plus7’ n = 10801) despite also having a similar number of expressed genes (‘Macrophages’ n = 2312 vs ‘Macrophages_Day31plus7’ n = 2219). Finally, NMP have a very low number of accessible peaks (n = 4071), in line with the low number of genes expressed in this cell state (n=1782) **(Fig. S4B)**.

### Transient activation of myeloid cells during the last phase of differentiation directs chromatin accessibility

For the macrophage differentiation phase, non-adherent cells at day 31 were collected and plated in fresh medium with cytokines for 7 days **(Fig. 3A)**. We analyzed the evolution of the cells (time points: day 31, day 31+1, day 31+4, day 31+7; **Fig. 3A top**), compared the effect of multiple cytokines on macrophage polarisation (cytokines: M-CSF, GM-CSF, GM-CSF+IL-34; **Fig. 3A top in red**) and tested the effect of using FBS vs. defined medium (media: RPMI+FBS, StemPro34; **Fig. 3A bottom**). We first performed an aggregated analysis of all experiments, annotating the cell types using LR from the fetal liver dataset^21^ **(Fig. 3A-B)**. In all experiments, we observed diverse cell types in distinct clusters that overlap regardless of the stimulation **(Fig. 3B)**. In addition, the distribution of the macrophages in the UMAP suggests that the major transcriptomic changes derive from differences in the time points and not the media composition (cytokine cocktails or base media used).

**Fig 3.**
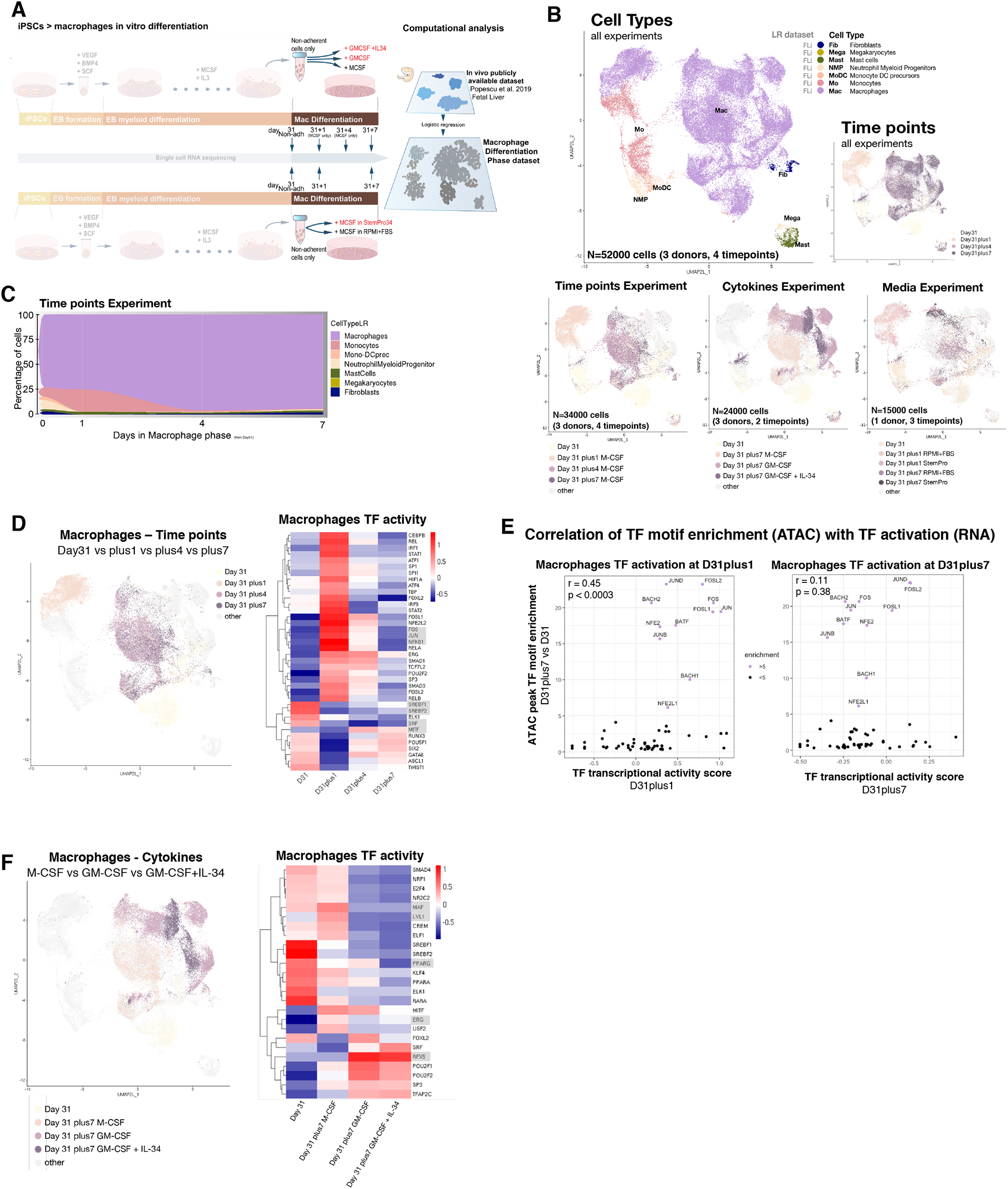
Evaluation of the macrophage phase. **A**, Schematic illustration of the *in vitro* differentiation protocol and cell-type annotation analysis with a focus on the time points of the macrophage phase (from day 31 to day 31 + 7). Alternative cytokine (top) and media (bottom) experiment highlighted in red. **B**, (top) UMAP projections of all the samples collected from the macrophage phase colored by cell type and time points. All experiments are pooled. (bottom) UMAP projections highlighting the samples included in each of the experiments colored by time point and condition. **C**, Stacked area plot of the cell-type percentages in each time point. Only samples from the time points experiment were included. **D**, (left) UMAP projection highlighting macrophages from the time points experiment (M-CSF only) and colored by time point. (right) Heatmap of the transcription factor activity scores calculated using DoRothEA across time points, relevant TF discussed in the text are highlighted in grey. **E**, TF motif enrichment values in macrophage ATAC open peaks at day 31+7 vs day 31 plotted against TF transcriptional activity score at day 31+1 (left) or day 31+7 (right). Pearson correlation’s r and significance are shown in the plots. **F**, (left) UMAP projection highlighting macrophages from the cytokines experiment and colored by time point and cytokine cocktail used. (right) Heatmap of the transcription factor activity scores calculated using DoRothEA+VIPER across time points and cytokines. iPSC, induced pluripotent stem cells; TF, transcription factor; ATAC, assay for transposase-accessible chromatin.

For the time points experiment, we analysed samples at day 31, day 31+1, day 31+4 and day 31+7 using M-CSF standard stimulation **(Fig. 3B)**. The non-adherent cells collected at day 31 from the EB myeloid differentiation phase were already mostly macrophages, alongside the main myeloid populations and a small subset of contaminating fibroblasts **(Fig. 3C)**. There was an enrichment in macrophages, representing a total of 94.1% of cells in the culture by day 31+7 **(Fig. 3C)**. This proportion is consistent with the CD14/CD64 surface protein levels obtained by FACS **(Fig. S5A)**. At the transcriptional level, there are broad differences between the macrophages of day 31 and day 31+1, as well as between day 31+1 and day 31+4/7, while macrophages from day 31+4 and day 31+7 completely overlap.

We analysed the major changes in the transcriptome of macrophages cultured in M-CSF over time by looking at TF activities^29,37^ **(Supplementary Table 6)**. From day 31 to day 31+1, we observed a transient but clear immune activation, as shown by increased activity of TFs such as JUN, FOS and NFKB1 **(Fig. 3D)**. This transcriptional activation returns to basal levels during day 31+4 and day 31+7. Despite being globally similar transcriptomically,, we observed few but relevant differences in TF activation between day 31 and day 31+7, including upregulation of MITF and downregulation of SRF, which regulate phagocytosis^38,39^, in addition to the downregulation of SREBF1 and SREBF2, involved in lipid metabolism and macrophage polarization^40,41^.

To assess whether the transient immune activation on day 31+1 affected chromatin structure, we analysed chromatin accessibility. We found that 113 TF motifs are significantly enriched in day 31+7 ATAC peaks, compared to day 31, and we obtained TF activity scores for 55 of these **(Fig. 3E, Supplementary Table 6)**. The top 11 enriched motifs (enrichment score >5) at day 31+7 correspond to TFs that were transcriptionally activated at day 31+1 but were no longer activated at day 31+7 **(Fig. 3E)**. Indeed, the global TF motif enrichment profile at day 31+7 is more correlated to TF activities at day 31+1 (Pearson correlation of TF activation at day 31+1: r = 0.45, p < 0.0003) than to activities at day 31+7 (Pearson correlation of TF activation at day 31+7: r = 0.11, p = 0.38) **(Fig. 3E)**. In short, this means activity on day 31+1 makes lasting changes on the chromatin landscape that are maintained until at least day 7 and may have transcriptomic consequences on future activations. Thus, this suggests the earlier macrophages represent a more naïve cell state, amenable to further reprogramming in response to polarization cues, which could have distinct applications.

Activated macrophages can be classified as M1 or M2 depending on whether they kickstart inflammation or resolve it, and the cytokines M-CSF and GM-CSF have been classically used to induce M2 and M1 primed phenotypes in macrophages, respectively^42^. We found that specific TFs, such as MAF, ERG and LYL1, had reduced activity scores in GM-CSF vs. M-CSF macrophages **(Fig. 3F, Supplementary Table 7)**. In contrast, RFX5, which is involved in MHC-II promoter activation, had increased activation in GM-CSF vs. M-CSF macrophages^32^. Despite GM-CSF being largely known to promote PPARG activation^35^, we found such an effect is reversed by the presence of IL-34 **(Fig. 3F)**. Of note, IL-34 is essential for the development of microglia from embryonic myeloid precursors^43^, and GM-CSF + IL34 induces the microglial phenotype on monocytes *in vitro*^*44*^. Both the knockdown and pharmacological antagonism of PPARG promotes LPS□stimulated transition from the M1 to the M2 phenotype in primary microglia, with the concomitant upregulation of markers such as CD206, TGFb and IL-4^45^.

Finally, the current iPSC-to-macrophage protocol uses chemically defined media with the exception of the last phase. We hypothesized that the inflammatory stimulation observed at day 1 of this phase is caused by the presence of FBS. Therefore, we tested whether using a defined medium (StemPro34 serum-free media, SP-SFM) at this step would reduce the observed activation. TF activity analysis showed SP-SFM induced similar activation signals. However, one notable difference was the maintained activity of the SREBF1 and SREBF2 TFs, which link lipid metabolism to the inflammatory response in macrophages **(Fig. S5B, Supplementary Table 7)**. This indicates that the composition of the medium affects macrophage metabolism and function, which needs to be taken into consideration during data interpretation^40,41^.

### iPSC-derived EB stimulation with GM-CSF and FLT3L produces type 2 conventional dendritic cells (cDC2)

We have shown that myeloid differentiation from iPSC produces a wide range of myeloid cells during differentiation and that the differentiation medium affects the regulatory programs that regulate cell identity. Conventional dendritic cells (cDC) present antigens to T cells and act as messengers between innate and adaptive immunity^46^. Protocols to induce DC differentiation *in vitro* are based on supplementing factors, including GM-CSF and FLT3L, that act cooperatively on cell precursors to drive cDC generation^47,48^. Based on that, we used GM-CSF and FLT3L (instead of M-CSF + IL3) in the EB myeloid differentiation phase and GM-CSF + IL4 (instead of M-CSF) in the last phase of differentiation on non-adherent cells from day 31 **(Fig. 4A)**. We annotated cells using LR classifiers that were trained on gastrulation^22^, embryonic yolk sac^6^ as well as fetal liver and thymus^20,21^ datasets **(Fig. 4A, Fig. S6, Supplementary Table 8)**.

**Fig 4.**
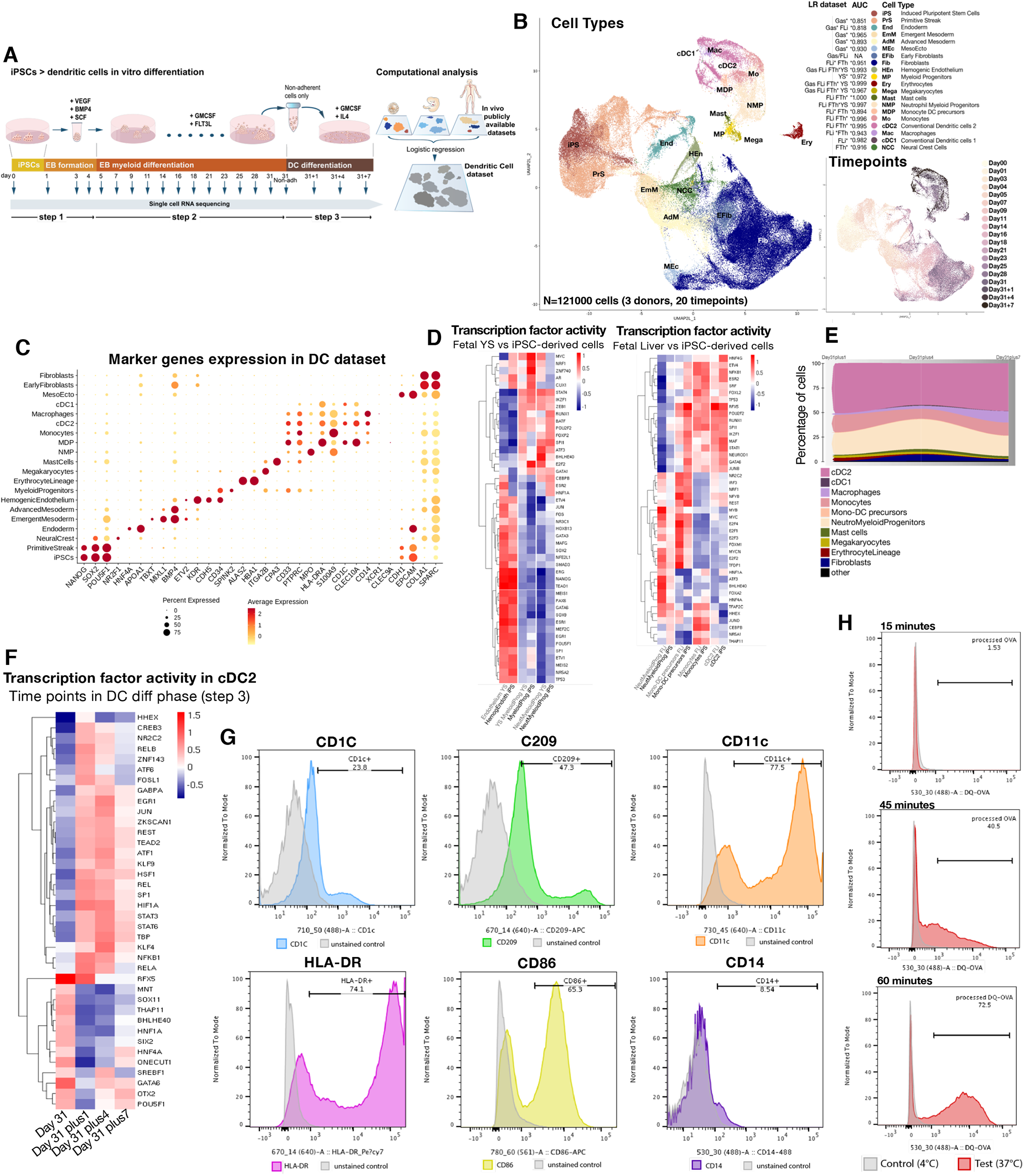
Modification of differentiation cytokines produces dendritic cells. **A**, (left) Schematic illustration of the *in vitro* differentiation protocol from iPSC to dendritic cells highlighting the time points when samples were collected for scRNAseq profiling. The protocol is analogous to the one used for differentiating iPSC into macrophages but employs distinct cytokines (GM-CSF and FLT3L for the EB myeloid differentiation phase and GM-CSF and IL-4 for the DC differentiation phase). (right) Diagram summarising the computational workflow used for cell-type annotation of the scRNAseq data generated with this protocol. **B**, UMAP projections of the scRNAseq data (n = 121,000) from 3 donors labeled by cell type (top) and time point (bottom). For each cell-type annotation we report, the *in vivo* datasets supporting the annotation and the mean area under the curve (AUC) for the logistic regression models trained on such datasets. **C**, Dot plot showing the average expression of canonical marker genes for each of the identified cell populations. **D**, Transcription factor activities computed with DoRothEA for the identified cell types present at day 21 of the *in vitro* differentiation protocol and matched cell types in the *in vivo* yolk sac dataset^6^ and fetal liver, skin and kidney dataset^21^, relevant TF discussed in the text are highlighted in grey. **E**, Stacked area plot showing the proportions of the major cell types produced by the differentiation protocol over the course of the last 7 days (Day 31+1 to Day 31+7). **F**, Transcription factor activities computed with DoRothEA for cDC2 identified at the last 4 time points of the differentiation protocol (step 3). **G**, Flow cytometry histograms showing the protein levels of cDC2 marker genes and CD14 as a negative marker in non-adherent cells at the end of the DC differentiation phase (day 31 + 7), matched unstained controls are shown in grey. **H**, Flow cytometry histograms for BODIPY™ FL DQ-ovalbumin processing by non-adherent cells at the end of the DC differentiation phase (day 31+7) incubated for 15, 45 and 60 minutes at 37°C. Matched samples were kept at 4°C for the same incubation periods and are shown in grey in each plot.

GM-CSF/FLT3L and M-CSF/IL3 stimulation induced the same cell types in the myeloid and stromal cell compartments **(Fig. 4B, Fig.1C)**, except that the modified protocol produced cells with a transcriptomic profile resembling cDC2 **(Fig. 4B, Fig. S6)**. Marker gene expression analysis further supported the cell type annotations, and specifically, cDC2 expressed bonafide cell-type markers (e.g., *HLA-DR, CD1C, CLEC10A*)^46^ **(Fig. 4C)**. A small subset of cells were assigned as cDC1 using LR but they do not express canonical cDC1 markers (e.g., *CLEC9A, XCR1*, **Fig. 4B-C, Fig. S6)**. The cell population dynamics were also similar between macrophage and DC protocols, with primitive streak, mesoderm and early fibroblasts being transient and the rest still present at the latest time point **(Fig. S7)**. As in the macrophage protocol, most myeloid populations arise simultaneously **(Fig. S7)**.

Cell types that arise during myelopoiesis using the DC protocol activate similar TFs *in vitro* and *in vivo* when compared to YS and fetal liver counterparts **(Fig. 4D)**, similar to what was observed using the macrophage protocol **(Fig, 2F-G)**. Specifically, the *in vitro* cDC2 activate TF networks relevant for *in vivo* cDC2 identity such as PU.1 (SPI1 gene)^49^ and KLF4^50^ **(Fig. 4D)**. We also observed increased RFX5 activity, which regulates MHC II gene expression^32^. A recent study postulated a role for CEBPB in the control of DC maturation and later stages of DC commitment^51^. Our results show reduced CEBPB activity in cDC2 cells compared to monocytes (*in vivo* and *in vitro*), which indicates that the *in vitro* phenotype shares features with a functionally mature DC subset characterized by upregulation of costimulatory and MHC class II molecules **(Fig. 4D)**.

We then evaluated the last phase of differentiation, following the non-adherent cells produced during EB myeloid differentiation until after the DC differentiation phase. A mean of 47.5% (standard deviation = 3.67) of the cells produced in the three time points analysed were cDC2 cells **(Fig. 4E)**, thus this differentiation is less efficient than the macrophage protocol, where macrophages represent 94.1% of cells by day 31 + 7 **(Fig. 3C)**. In contrast to what is observed with macrophages, the proportion of cDC2 remains stable **(Fig. 3C, Fig. 4E)**. The *in vitro* activation of cDC2 and macrophages induce shared regulatory programs, including activation of NFKB1 or JUN on day 1 of this phase **(Fig. 3C, Fig. 4F, Supplementary Table 9)**. Interestingly, the particular profile of TF networks induced in cDC2 on that day (including JUN, REL, SP1 and HIF1A) does not fully return to basal levels by day 31+7 **(Fig. 4F, Supplementary Table 9)**. This is in contrast to what is observed in the macrophage differentiation phase **(Fig. 3D, Supplementary Table 5)**.

Finally, in order to confirm the cDC2 phenotype of the iPSC-derived cells, we checked the expression levels of cDC2 canonical surface proteins using FACS and functionally interrogated cDC2 using an antigen processing assay (DQ-OVA). We observed positive cells for cDC2 markers (CD1C, CD209, CD11c, HLA-DR, CD86) and markedly low levels of CD14, which is a marker for macrophages, thus validating the cDC2 phenotype^52^ **(Fig. 4G)**. The cross-presentation capacity, as well as the kinetics of antigen uptake and proteolytic degradation, differ in human DC subtypes^53^. Functionally, adult monocyte-derived DC have higher antigen uptake and processing at earlier time points (15 to 30 min), whereas cDC processing capacity peaks at 60 min^54^. To look at the antigen processing activity in our iPSC-derived cDC2, we measured BODIPY-conjugated DQ-OVA processing at several time points (15 min, 45 min and 60 min). iPSC-derived cDC2 showed no DQ-OVA processing at shorter time points (15 min) but did process OVA at longer time points (45 min and 60 min) **(Fig. 4H)**. These results reveal that our cDC2’s antigen processing behaviour resembles that of classical DC and reinforces the idea that a functionally mature cDC2-like cell can be recapitulated from iPSC under these conditions.

### Genes associated with immune phenotypes through GWAS shape the transcriptomic profile of iPSC-derived myeloid cells

Dysfunctional myeloid differentiation and signaling downstream of myeloid receptors lead to immune-related disorders^55^. To dissect potential myeloid contributions involved in these pathologies, we selected four genes linked to immune-related GWAS hits (i.e., *ICAM1, LSP1, PRKCB* and *ZEB2*) based on the existing literature. The *ICAM1* and *LSP1* loci contain SNPs linked to autoimmune inflammatory diseases by GWAS (https://www.ebi.ac.uk/gwas/home) and interact with each other^56^. *PRKCB* is a protein kinase associated with inflammatory diseases and blood cell counts in GWAS data (https://www.ebi.ac.uk/gwas/home) and is involved in myeloid DC differentiation^57^. Finally, *ZEB2* is found in GWAS for blood phenotypes and regulates hematopoiesis in mice^58^ and the cell fate decisions of DC^59^. To study their involvement in myeloid differentiation and identity, we generated knock-out (KO) iPSC lines using CRISPR/Cas9 in one of our cell lines (KOLF-iPSC).

CRISPR/Cas9 KO iPSC lines were differentiated into macrophages and DC alongside isogenic WT lines. Single-cell transcriptomic analyses were performed at day 0 (iPSC stage) and day 31 (EB myeloid differentiation) **(Fig. 5A-B)**. Though we observed all cell populations in all conditions **(Fig. 5C-D)**, some of the KOs affected the cell type proportions. As expected, knocking out *ZEB2* reduced the proportion of myeloid cells to 5.5%, versus 65.6% in WT lines, in the macrophage protocol (12-fold decrease) and 2.1%, versus 20.4% in WT lines, in the DC protocol (10-fold decrease) **(Fig. S8A, Supplementary Table 10)**. On the contrary, *PRKCB* KO increased the proportion of myeloid cells but only during the DC differentiation protocol (20.4% of cells in WT vs 78% in *PRKCB* KO, 4-fold increase) **(Fig.S8A, Supplementary Table 10)**. The other KOs did not seem to influence the ratio of myeloid cells

**Fig 5.**
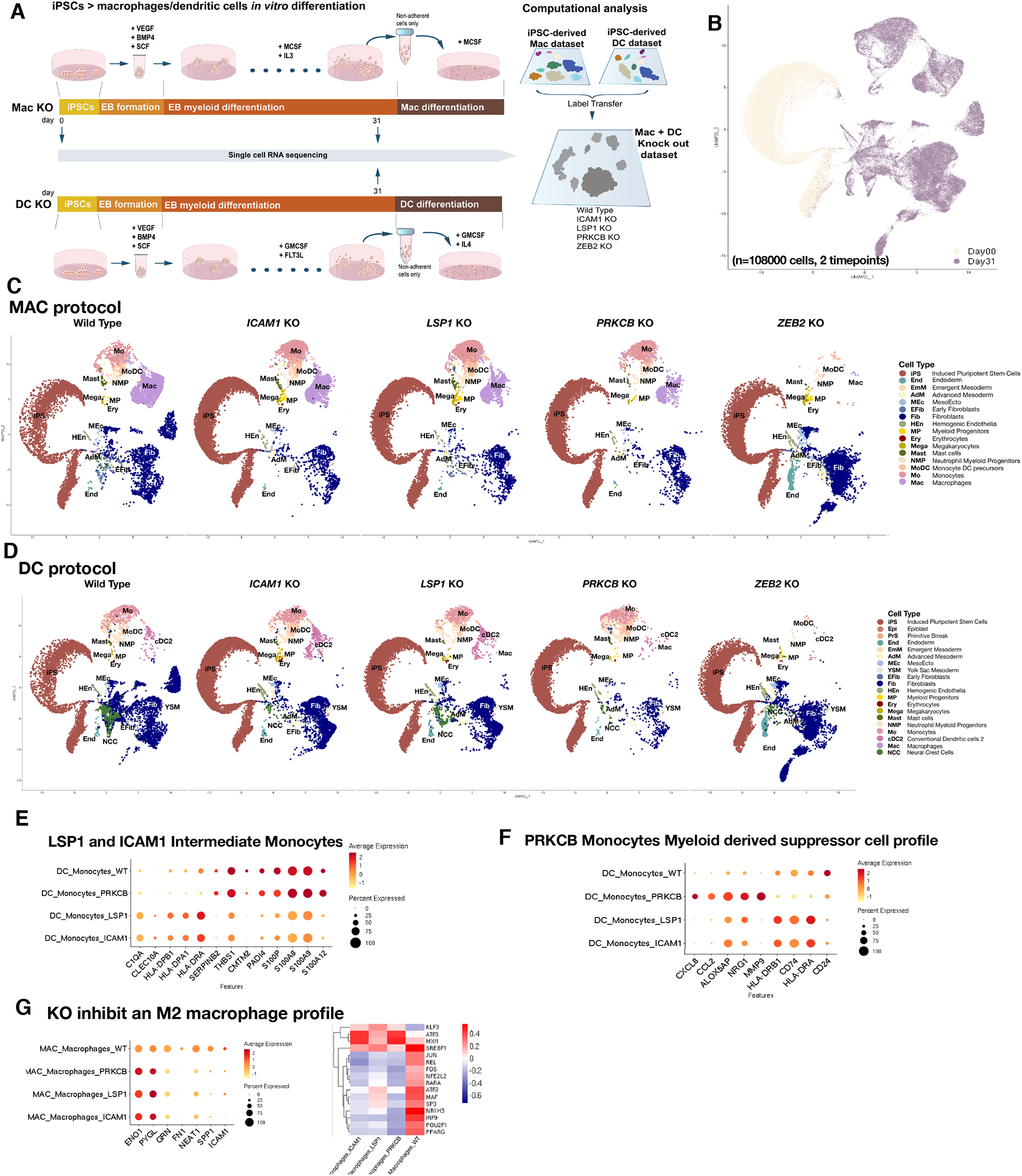
Effect on macrophage differentiation of *ICAM1, LSP1, PRKCB* and *ZEB2* KO. **A**, (left) Schematic illustration of the *in vitro* differentiation protocols from iPSC to macrophages (top) or dendritic cells (DC, bottom) used to evaluate the effects of *ICAM1, LSP1, PRKCB* or *ZEB2* knockouts (KO). Samples were collected at day 0 and day 31 of the protocols and profiled with scRNAseq. (right) Diagram summarising the computational workflow used for cell-type annotation of the scRNAseq data generated with these protocols. Briefly, cell-type annotations were transferred from scRNAseq data of the macrophages (Discovery dataset) and DC protocols described in the previous sections. **B**, UMAP projections of scRNAseq data (n = 108,000) from both KO protocols labeled by time point. **C**, UMAP projections of scRNAseq data generated from the iPSC-to-macrophages KO protocol (one UMAP per KO plus wild type) coloured by cell type. **D**, UMAP projections of scRNAseq data generated from the iPSC-to-DC KO protocol (one UMAP per KO plus wild type) coloured by cell type. **E**, Dot plot showing the average expression of intermediate monocyte–associated genes in the monocytes produced by each KO and the wild type in the iPSC-to-DC protocol. **F**, Dot plot showing the average expression of genes associated to myeloid-derived suppressor cells in the monocytes produced by each KO and the wild type in the iPSC-to-DC protocol. **G**, (left) Dot plot showing the average expression of M2-associated genes in the macrophages produced by each KO and the wild type in the iPSC-to-macrophages protocol. (right) Transcription factor activities computed with DoRothEA for macrophages produced by each KO and the wild type.

At the transcriptomic level, the phenotypes were different between KO lines. Monocytes from the *LSP1* and *ICAM1* KOs had a transcriptomic profile that coincided with that of intermediate monocytes, characterised by the upregulation of the HLA genes and downregulation of the S100 gene family^60^ **(Fig. 5E)**. Interestingly, this profile is not observed in the KO monocytes produced by the macrophage differentiation protocol **(Fig. S8B)**. An increased population of intermediate monocytes has been described in many autoimmune diseases such as active Crohn’s disease^61^ and rheumatoid arthritis^62^. Monocytes from the *PRKCB* KO cell line after the DC differentiation protocol had a myeloid-derived suppressor profile, including low expression of *HLA-DR* and *CD74* with high levels of *CCL2* and *MMP9*^*63–66*^ **(Fig. 5F)**. This is consistent with the observation that myeloid-derived suppressor cells have decreased levels of *PRKCB*, which dampens DC differentiation and function *in vivo*^*57*^.

Macrophages generated from iPSC deficient in *PRKCB, LSP1* or *ICAM1* exhibited a mixed anti-inflammatory and anti-fibrotic phenotype. *PRKCB, LSP1* or *ICAM1* KO macrophages upregulated the suppressors of the NFkB-dependent inflammatory pathway KLF3^67^ and ATF3^68^ **(Fig. 5G)**. They also decreased the expression of genes linked to an M2 profibrotic macrophage phenotype (e.g., *FN1, GRN* and *SPP1*, **Supplementary Table 11**), as well as decreased activity of M2-promoting transcription factors (e.g., MAF^33^ and PPARG^69^, **Fig. 5G**). *PRKCB, LSP1* and *ICAM1* are connected to each other, as *ICAM1* is downregulated in *PRKCB* and *LSP1* KOs **(Supplementary Table 11)**. These results suggest that *PRKCB, LSP1* and *ICAM1* are part of a regulatory network that fine tunes macrophage phenotype and represses tissue healing both by promoting REL/p65-mediated inflammation and controlling the expression of profibrotic M2 genes. Notably, silencing *PRKCB, LSP1* and *ICAM1* generates a macrophage population with a mixed phenotype characterized by the inhibition of M2 tissue remodelling programs **(Fig. 5G)** and the suppression of the NFKB pathway. In this respect, such broad defects in macrophage polarization could impair proper resolution of inflammation, leading to *in vivo* autoimmune inflammatory disorders^70^.

Altogether, we have shown that iPSC-differentiation protocols are powerful tools to interrogate specific genes mediating early hematopoiesis (e.g., *ZEB2*) as well as monocyte and macrophage function (e.g., *ICAM1, LSP1* and *PRKCB*). We found changes in the inflammatory potential of the myeloid cells lacking expression of genes associated with autoimmunity, coinciding with the expected phenotype.

## Discussion

The full characterisation and assessment of the robustness, accuracy and efficiency of *in vitro* protocols is essential to utilising them as models for disease as well as leveraging them in the search for novel therapeutic targets. Myeloid cells have a central role in immunity and are involved in major inflammatory and autoimmune disorders. The establishment of robust experimental protocols to generate macrophages that are easy-to-replicate and amenable to scaling up is paramount to studying human macrophage ontogeny, genetics and function in health and disease. Here, we profiled more than 400k single cells across a commonly used, straightforward differentiation process from human iPSC to myelopoiesis to terminally differentiated macrophages. We reconstructed, using cell trajectories, the *in vitro* sequence of events, which parallel fetal hematopoiesis prior to the establishment of HSC. Moreover, this protocol is a valuable resource in studying multiple myeloid populations, including erythrocytes, megakaryocytes and mast cells that spontaneously arise, as well as DC, that can be induced by adjusting the composition of the media. Finally, we used this model to interrogate the functional effect of genes associated with inflammatory and autoimmune disorders and interpreted the results in relation to their *in vivo* counterparts.

To quantitatively assess the accuracy of our *in vitro* models, we used machine learning tools. We built logistic regression models trained on scRNAseq data from developmental atlases mapping the formation of the immune system and projected it onto the *in vitro* datasets. The computational framework we have established in the work could be adapted to annotate cells arising from multiple iPSC differentiation protocols. Following this strategy, we found that the initial phases of iPSC–macrophage differentiation faithfully recapitulate YS fetal hematopoiesis and generate fetal-like *FOLR2+* macrophages. The lack of an HSC cluster in our data, the activation of master regulator RUNX1 in the endothelial-to-hematopoietic transition (EHT)^71^, and the lack of expression of HOXA genes in the myeloid progenitors^24^ all suggest our protocols recapitulate YS differentiation prior to the establishment of definitive hematopoiesis. Thus, we propose this model as a unique system for interrogating the early stages of hematopoietic differentiation in humans, which are largely unexplored.

Our study shows that iPSC–macrophage differentiation generates a wide range of myeloid cells and presents a detailed list of TFs that mediate the generation of distinct myeloid progenitors *in vitro*. We also observed that supplementation of the culture media with factors, particularly GM-CSF and FLT3L, drives iPSC-derived myeloid progenitors into cell types that express markers of cDC2^46^. This result uncovers the potential of these cytokines to promote DC-like cell identity, either directly or through monocytes. The protocol for DC generation had a lower efficiency than the MAC-producing protocol, which is consistent with the main differentiation of YS myeloid progenitors towards macrophages^6,19–22^. Interestingly, we found erythrocytes in our data following a decline in RUNX1 and SPI1 TF activity^72^ and a rise in GATA1 activation^73^. The *in vitro* differentiation of erythroid-lineage cells from iPSC has relevant biomedical implications for their use as disease models in the study and treatment of anemias^74^. Monocytes are also generated during differentiation but our trajectory analysis indicates that macrophages are derived from a myeloid intermediate and bypass the monocyte stage. It is tempting to speculate that isolated monocytes from this protocol can also be differentiated into macrophages and polarised to specific functions in the presence of tissue-specific environmental signals^75,76^. Future work should evaluate if the origin of macrophages imprints on their function.

iPSC-derived macrophages stimulated with M-CSF compared to unstimulated cells show greater global chromatin accessibility that is not a reflection of an increased number of genes expressed, indicating epigenetic rewiring. Epigenetic states in macrophages are instrumental in the generation of functional and phenotypic diversity^76–78^. Here, we demonstrated that exposure to M-CSF generates macrophages with a transcriptomic profile similar to their unstimulated counterparts, but an increase of the chromatin accessibility points to a reprogramming of the epigenetic landscape. We also identified multiple features resembling YS myelopoiesis in our iPSC system. Nonetheless, despite the YS origin of Kupffer cells (KC)^76,79^ the LR classifier trained specifically on fetal liver KC does not capture any cell from the *in vitro* dataset, indicating that the strong tissue-resident signature of these cells is not recapitulated using this protocol. In particular, we did not observe activation of KC-determining TF LXRa, RBPJ or SMAD4 probably due to a lack of liver-derived signals such as Notch ligand DLL4 essential for their induction^80^. All considered, the iPSCs protocol to derive certain macrophage subtypes, combined with high resolution single cell analysis provides the unprecedented possibility to directly interrogate the extent of macrophage’s intrinsic plasticity, which remains a matter of debate^81^. A marked enrichment of the macrophage population was observed during the macrophage differentiation phase, pointing to the appropriateness of using single-cell approaches to unravel cell heterogeneity in the initial phases (day 31 and day 31+1) and bulk technologies at later time points (day 31+4 and onwards). Our combined transcriptome/ATAC analysis shows the potential of macrophages, among non-adherent cells in the EB myeloid differentiation phase, to be truly naïve cells early on, whereas 24h into the macrophage differentiation phase, there are M-CSF-induced and reversibly activated macrophages.

Finally, we leveraged this model to experimentally evaluate genes linked to immune-related disorders by GWAS. Interestingly, CRISPR-Cas9-mediated knockout of GWAS hits (*PRKCB, LSP1* and *ICAM1*) in iPSC-derived macrophages revealed their role in the regulation of both inflammatory signaling and extracellular matrix (ECM) deposition. Fibrosis constitutes a pathological feature of most chronic inflammatory diseases including the ones featured in our study^82,83^, and our results open an avenue of therapeutic intervention in these disorders. In line with this, we show that macrophages obtained through this protocol recapitulate TAM states in the liver tumour microenvironment. While fetal-like FOLR2+ TAMs are more similar to stimulated macrophages after day 4 and 7, SPP1+ TAMs resemble the macrophage state at day 1. This suggests these cells could also be a faithful model to recapitulate macrophage subtypes in the tumour microenvironment.

In conclusion, we have defined a comprehensive map of cells and molecular programs that underlie iPSC–macrophage differentiation in a dish. Macrophages play an important role for immunity in health and disease, and represent key cellular targets for immunotherapy. Our study shows the potential of deeply characterizing differentiation protocols at the single-cell level and demonstrates that it is a valuable model for interrogating the very early stages of hematopoietic formation that have been largely unexplored so far.

## Material and Methods

### Human induced pluripotent stem cell lines

All iPSC lines used in the study were generated by the HIPSCI project. Details on their generation are available at http://www.hipsci.org. Briefly, we used kolf_2, yemz_1 and vass_1 in the Discovery and DC datasets, and we added ceik_1, eesb_1 and wegi_1 for the Validation datasets. All cells in the knockout dataset are derived from kolf_2 as a parental line. All HIPSCI samples were collected from consenting research volunteers recruited from the NIHR Cambridge BioResource (http://www.cambridgebioresource.org.uk), initially under existing ethics rules for iPSC derivation (Regional Ethics Committee (REC) reference 09/H0304/77, v.2, 4 January 2013), with later samples collected under a revised consent (REC reference 09/H0304/77, v.3, 15 March 2013).

### *In vitro* differentiation to macrophages and dendritic cells

We used an adaptation of the van Wilgenbrug et al. protocol^15^. Feeder-free human iPSC were cultured in E8. For the embryoid body (EB) formation, step1, a single-cell suspension of hiPSC was plated in 100 µl of EB medium – E8 + SCF (20 ng/ml) + VEGF (50 ng/ml) + BMP-4 (50 ng/ml) + ROCK inhibitor (10 µM) – at a density of 10,000 cells per well in round bottom low-attachment 96 well plates. After 2 days, we changed half the media (50 µl) and replaced it with fresh EB media. At day 4, EB myeloid differentiation started, step 2, when EBs were plated in gelatin-coated 6-well plates at a density of 8–10 EBs per well in EB-Mac medium – StemPro-34 + M-CSF (100 ng/ml) and + IL-3 (25 ng/ml). The EB-Mac medium was changed every 4 to 5 days. At day 31, step 3, non-adherent cells were collected by centrifugation with the medium change and cultured in 10 cm tissue culture plates for 7 days in macrophage differentiation medium – RPMI + 10% heat-inactivated FBS + M-CSF (100 ng/ml).

Alternative macrophage differentiation media were used in the macrophage differentiation phase, step 3. For the cytokines experiment **(Fig. 3B-D)**, we used RPMI + 10% heat-inactivated FBS + GM-CSF (50 ng/ml) and RPMI + 10% heat-inactivated FBS + GM-CSF (10 ng/ml) + IL-34 (100 ng/ml). For the media experiment **(Fig. 3B, Fig. S5B)**, fully defined medium – StemPro-34 + M-CSF (100 ng/ml) -was used.

Step 1 is shared between macrophages and DCs, while different cytokines are used in step 2 and 3. For DC differentiation EBs at day 4 are plated with EB-DC media – StemPro-34 + GM-CSF (50 ng/ml) + FLT3L (100 ng/ml) in the same types of plates and density as the macrophage protocol. At day 31, step 3, non-adherent cells were collected and plated in 10 cm tissue culture plates in DC differentiation medium – RPMI + 10% heat-inactivated FBS + GM-CSF (50 ng/ml) + IL-4 (100 ng/ml).

### 10x Genomics Chromium GEMs sample preparation and sequencing

Single-cell transcriptomic analysis on iPSC-to-macrophage differentiation was performed in 3 iPSC lines for the Discovery dataset and 6 hiPSC lines for the Validation dataset. One 6-well well per line was collected using TrypLE at 20 timepoints in the Discovery dataset, and 2 6-well wells per line at 7 timepoints in the Validation dataset, between Day 0 and Day 38 (Day 31 EBs plus 7 days of the macrophage differentiation phase). At every collection day, cells of each well were merged, counted, passed through 40 µM filters and resuspended in DPBS + 0.4% BSA. Cell suspensions were processed using the Chromium Single Cell 3’ kit (v2 for Discovery, v3 for Validation), aiming at recovering from 3000 to 10000 cells. Library preparation was carried out according to the manufacturer’s instructions. Libraries were sequenced, aiming at a minimum coverage of 50000 raw reads per cell, on the Illumina HiSeq 4000 (Discovery) or Novaseq 6000 (Validation) using the sequencing formats; read 1: 26 cycles; i7 index: 8 cycles, i5 index: 0 cycles; read 2: 98 cycles (3’ kit v2) or read 1: 28 cycles; i7 index: 8 cycles, i5 index: 0 cycles; read 2: 91 cycles (3’ kit v3).

Sample preparation and sequencing for the DC datasets was performed as described for the Discovery dataset. The Knockout dataset samples were processed as described for the validation dataset but only for 2 time points (i.e., Day 0 and Day 31).

Single-cell ATAC analysis was performed in a subset of the single-cell suspensions for 6 of the time points of the Validation dataset described above. Single-nuclei suspensions were obtained and processed according to the manufacturer’s instructions using Chromium Single Cell ATAC v1.0, aiming for 10000 nuclei per sample. Library preparation was carried out according to the manufacturer’s protocol and sequenced on Illumina NovaSeq 6000, aiming for 20000 fragments per cell using the sequencing formats; read 1: 50 cycles; i7 index: 8 cycles, i5 index:16 cycles; read 2: 50 cycles.

### Single-cell RNA seq computational analysis

Cell Ranger (v3.1.0), mapping to GRCh38 (v3.0.0), was used to filter out empty droplets using default values. Cells were further filtered out for the number of genes (<200) and percentage of mitochondrial RNA (>8.5%) using Seurat (https://satijalab.org/seurat/v3.2.2). All cells identified as doublets using SoupOrCell^84^, DoubletDetection (http://doi.org/10.5281/zenodo.2678041) and Scrublet^85^ were discarded. Cell genotype calling was performed using SoupOrCell^84^. All datasets were normalized using sctransform in Seurat^86^, and UMI counts, mitochondrial RNA and cell cycle variables were regressed out by cell line. Multiple hiPSC lines were integrated using Seurat’s anchor-based method^23^. After PCA dimensionality reduction and louvain clustering^87^, datasets were further analysed as described below.

### Single-cell ATAC seq computational analysis

Cell Ranger ATAC pipeline (v1.2.0), mapping to GRCh38 (v3.0.0), was used for read filtering and barcode cell calling. Peaks were re-called using cellatac, an in-house implementation of Cusanovich’s approach^88^ (https://github.com/cellgeni/cellatac)^89^. Peak and cell filtering were performed using cellatac and Signac (https://satijalab.org/signac/version1.1.1), as described in Fig. S1B. Normalization and dimensionality reduction were performed using term frequency - inverse document frequency (TF-IDF) and Singular Value Decomposition (SVD), respectively. SLM from Seurat was used for clustering. TF motif analysis was performed using Signac and JASPAR 2020^90^ motifs database.

### Cell-type annotation of RNA and ATAC datasets

Both the Discovery and DC datasets were annotated using logistic regression (LR) models built on publicly available single-cell transcriptomic datasets. The LR prediction models used at each step were built based on a general linear model function and a 10-fold cross-validation. Briefly, public raw data (Cell Ranger output when available or processed matrices otherwise) was downloaded and re-processed as described for the Discovery and DC datasets. Then, public datasets were split in training (70%) and test (30%) sets, ensuring these proportions were accounted for each cell type. Then we generated LR models to classify each cell type for each gene in the training partition of the *in vivo* dataset using normalised data. A ranked gene list based on the area under the curve (AUC) of each gene was produced for each cell type. The optimal number of genes to build the final LR classifier was chosen by building models on the training set and calculating the AUC of the prediction on the test set. This was repeated with an increasing number of genes down the ranked list described above. The number of genes that produced a model on the training set with the highest AUC when applied to the test set was then used to build the final model on the full *in vivo* dataset. This LR prediction model was then used to classify the cells in the *in vitro* dataset. Finally, the mean prediction probability per louvain cell cluster was calculated for all the LR models built, and each cluster was labeled based on the LR model with the highest mean prediction. As an estimate of the strength of the association between the annotation and the labeled clusters, we calculated the AUC of each annotated cell type based on the LR probabilities.

For the validation dataset, cell type annotations from the Discovery dataset were projected on the transcriptomic and ATAC validation datasets using Seurat’s anchor-based label transfer approach^23^.

### Trajectories analysis

Spliced/unspliced RNA expression matrices were generated using the command line tool from velocyto (http://velocyto.org/velocyto.py/tutorial). scVelo was used for trajectory analysis based on RNA velocity and PAGA graph abstraction as described (https://scvelo.readthedocs.io/DynamicalModeling/). All analyses were performed on a per sample basis.

### Transcription factor activity analysis

Transcriptomic changes across trajectories and time points were studied based on transcription factor activities using DoRothEA and VIPER analysis^29^. DoRothEA v1.2.1 (https://saezlab.github.io/dorothea) required Seurat v4.0.2 (https://satijalab.org/seurat/). Both *in vitro* and *in vivo* datasets were subset based on connected cell types according to the trajectory analysis. Normalised data was scaled within each subset, and TF activity scores were computed for each cell for 271 TFs with high-confidence target-gene annotation (A, B and C confidence levels, https://saezlab.github.io/dorothea/). Heatmaps for *in vitro* vs *in vivo* comparison were produced by selecting the top 50 most variable TFs in each dataset, and results were merged and plotted using pheatmap (https://www.rdocumentation.org/packages/pheatmap/versions/1.0.12).

### Marker protein and antigen processing DQ-OVA assay analysis by FACS

Macrophages and dendritic cells were detached from 10 cm plates using Lidocaine + EDTA for 5 min at 37°C, collected in DPBS and spun down at 300g for 3 min. Samples were then fixed with BD Cytofix buffer for 20 min at room temperature and washed with DPBS + 1% FBS. Staining with fluorescent-labeled primary antibodies was performed in the dark at room temperature for 30 min. After 2 washes with DPBS +1% FBS, cells were analysed by FACS in a BD LSR Fortessa II.

Dendritic cells were collected as described above and incubated with DQ-OVA in the dark at 4°C or 37°C for 15 min, 45 min and 60 min, as indicated. Cells were then washed with ice-cold DPBS +1% FBS and analysed by FACS as above.

### CRISPR-Cas9 knockout of human induced pluripotent stem cell lines

Knockout iPSC lines were generated by substituting an asymmetrical exon with a Puromycin cassette and expanding those clones with a frame-shift indel in the remaining allele. A hSpCas9 and two small guide RNA expression vectors along with a template vector were used. The template vector harboured an EF1a-Puromycin cassette with two flanking 1.5 kb homology arms designed around the asymmetric exon of interest. For each knockout line, 2×106 iPSC single cells were transfected using the Amaxa Human Stem Cell Nucleofector® Kit 2 (Lonza) with 4 μg, 3 μg and 2 μg of each plasmid, respectively, and plated in 10 cm plates. After 72 h, cells were selected in 3 μg/mL Puromycin and colonies were expanded and genotyped. Lines confirmed to have the Puromycin cassette and the presence of a frame-shift indel by Sanger sequencing were selected for the experiments.

### Differential expression analysis of knockout iPSC-derived cell types

Transcriptomic alterations between cell types arising in WT and KO lines were assessed using differential expression from Seurat. Genes present in 10% of the cells and with a minimal log fold-change of 0.25 were selected for differential expression analysis of each cell type in each KO line vs their WT counterpart. Only genes with an FDR < 0.05 were considered as significantly differentially expressed.

## Acknowledgments

We thank the Cellular Genetics wet lab support team, Cellular Genetics IT team, Sanger Sequencing operations and Sanger Cytometry Core facility for their essential help. We thank the Gene Editing team for providing iPSC knock out lines. We would also like to thank Ruxandra Tesloianu and Luz Garcia-Alonso for their help setting up the scATAC-seq computational analysis, and Carmen Sancho-Serra for her assistance in the experiments. We thank Jana Eliasova for her help with figure design and Christina Usher for her edits in the text. This work was supported by the Open Targets consortium (OTAR026 project) and the Wellcome Sanger core funding (WT206194). The authors are grateful to the funders for their support and additional care given to their members during the COVID-19 pandemic. D.A-E thanks CERCA Programme/Generalitat de Catalunya and the Josep Carreras Foundation for institutional support. This study makes use of cell lines and data generated by the HipSci Consortium, funded by The Wellcome Trust and the MRC (Medical Research Council). For the purpose of Open Access, the author has applied a CC BY public copyright licence to any Author Accepted Manuscript version arising from this submission.

## Author contribution

C.A, D.G and R.V-T conceived and designed the experiments and analyses. C.A and M.P performed the experiments with contributions from C.S-S. A.K designed and performed the ATAC single cell sample processing. C.A analysed the data with contributions from V.L with the trajectories and knock out analyses, from J-E.P with the thymus dataset logistic regression and from B.W and D.T with the dendritic cells dataset. C.A, R.V-T and D.E-G interpreted the data and wrote the manuscript with contributions from V.L. D.G and R.V-T supervised the work. All authors read and approved the manuscript.

## Data availability

Datasets are being uploaded into ArrayExpress, and will be available through the web portal www.HiPImmuneatlas.org. All code used for data analysis is being made available from https://github.com/Ventolab/iPSCmyeloid.

## Supplementary figure legends

**Supplementary Figure 1. Computational workflow**

**A**, Computational processing and analysis of 10X Genomics Chromium single-cell GEMs 3’ RNA samples. **B**, Computational processing and analysis of 10X Genomics Chromium single-nuclei GEMs ATAC samples.

**Supplementary Figure 2. Logistic regression predictions from *in vivo* datasets for cell types in the Discovery dataset**

Discovery dataset UMAP projections showing the logistic regression prediction probabilities for models trained on each cell type present in publicly available single-cell transcriptomic datasets. Prediction probabilities built on: **A**, Human gastrulation embryo dataset, 2-3 post conceptional weeks (PCW)^22^, **B**, Human fetal liver, skin and kidney cells, 7-17 PCW^21^, **C**, Human fetal yolk sac 4-5 PCW^6^, **D**, Human fetal thymus and liver cells, 7-17 PCW^20^ and, **E**, Human fetal, 6-12 PCW, and decidual, adult, cells^19^.

**Supplementary Figure 3. Further characterisation of the Discovery dataset**

**A**, Discovery dataset UMAP projections showing scaled gene expression for the HOXA family of genes and the location of cells annotated as myeloid progenitors. **B**, UMAP projections of the yolk sac dataset^6^ coloured by the predicted probabilities by logistic regression of myelopoiesis cell types from the *in vitro* iPSC-derived Discovery dataset, (bottom) UMAP projections of the cell types described in the yolk sac study^6^ and cell types identified through logistic regression analysis. **C**. Heatmap showing the mean predicted probabilities by logistic regression of the cell types found in the yolk sac ^6^. In red are cell type clusters not described in the original yolk sac study^6^ and defined by the logistic regression results in B. **D**, Discovery dataset UMAP projections, first with the cell type annotations as reference and the rest are coloured by the predicted probabilities by logistic regression of each macrophage subtype found in the maternal–fetal interface^19^. **E**, Heatmap showing the mean predicted probabilities by logistic regression of the macrophage subtypes found in the maternal–fetal interface^19^ for each of the cell types in the Discovery scRNAseq dataset.

**Supplementary Figure 4. Additional supporting data for the trajectories analysis**

**A**, Heatmap of the percentage of cells distribution across all timepoints for each cell type in the Discovery dataset. **B**, Violin plots of the mean number of genes expressed per cell in each of the cell types across the main differentiation trajectories identified. Black dot = median number of expressed genes per cell type.

**Supplementary Figure 5. Additional supporting data for the macrophage phase**

**A**, FACS plots showing CD14 and CD64 surface protein expression levels for cells collected at the end of the differentiation, plus7, for the three donor iPSC lines used in the discovery dataset. **B**, (right) Macrophage phase UMAP projection highlighting macrophages from the media experiment and colored by time point and media composition. (left) Heatmap of the transcription factor activity scores calculated using DoRothEA across timepoints and media composition.

**Supplementary Figure 6. Logistic regression predictions from *in vivo* datasets for cell types in the DC dataset**

Dendritic cell dataset UMAP projections showing the logistic regression prediction probabilities for models trained on each cell type present in publicly available single-cell transcriptomic datasets. Prediction probabilities built on: **A**, Human gastrulation embryo dataset, 2-3 post conceptional weeks (PCW)^22^, **B**, Human fetal liver, skin and kidney cells, 7-17 PCW^21^, C, Human fetal yolk sac, 4-5 PCW^6^ and **D**, Human fetal thymus and liver cells, 7-17 PCW^20^.

**Supplementary Figure 7. Dendritic cells protocol**

**A**, Heatmap of the percentage of cells distribution across all time points for each cell type in the DC dataset.

**Supplementary Figure 8. Knock-out cell types’ transcriptomic profiles**

**A**, Stacked bar plots of the proportions of each cell type for each KO gene and WT lines on day 31 cells for the macrophage (left) and dendritic cells (right) protocols. **B**, DotPlot with scaled gene expression levels in monocytes from WT and knock-out lines differentiated with the macrophage protocol. Genes shown are characteristic of intermediate monocytes and were significantly dysregulated in LSP1 and ICAM1 monocytes produced in the DC protocol.

