## Supplementary Figures for "Robust temporal map of human *in vitro* myelopoiesis using single-cell genomics"

**Supplementary Figure 1.**

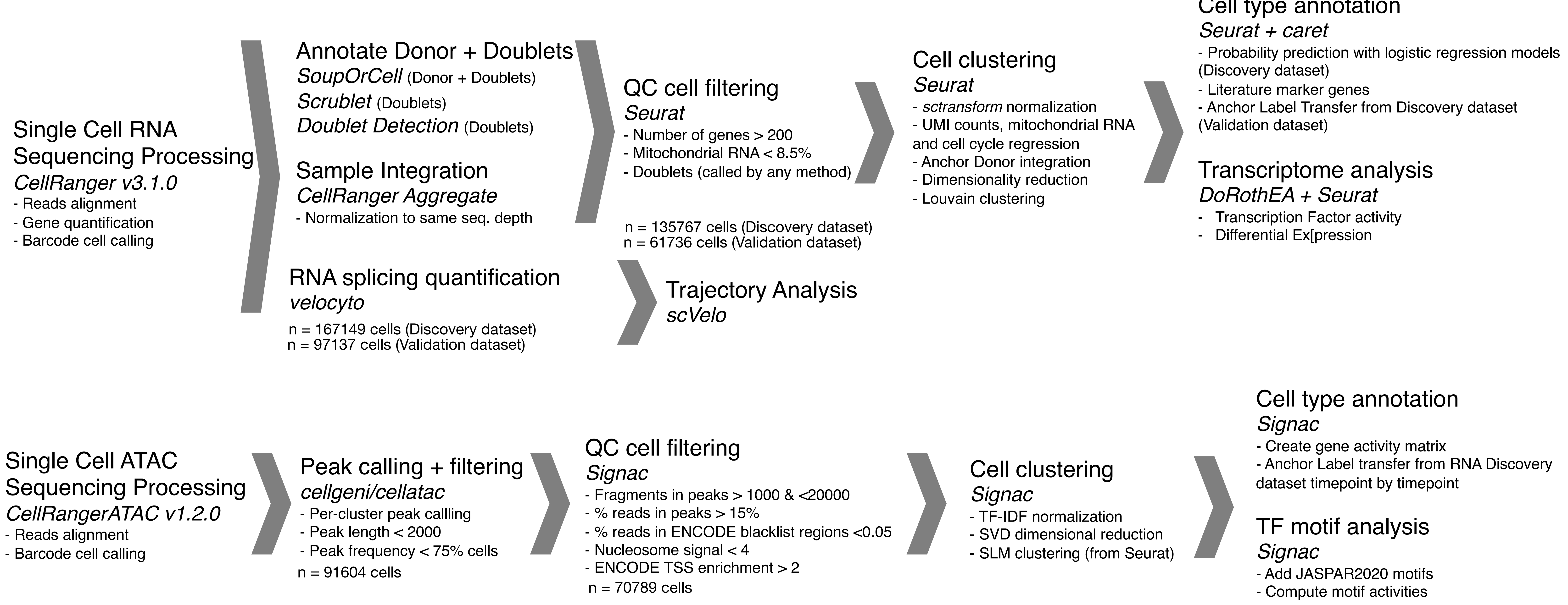

Supplementary Figure 2.

**A** **GASTRULATION dataset cell types**

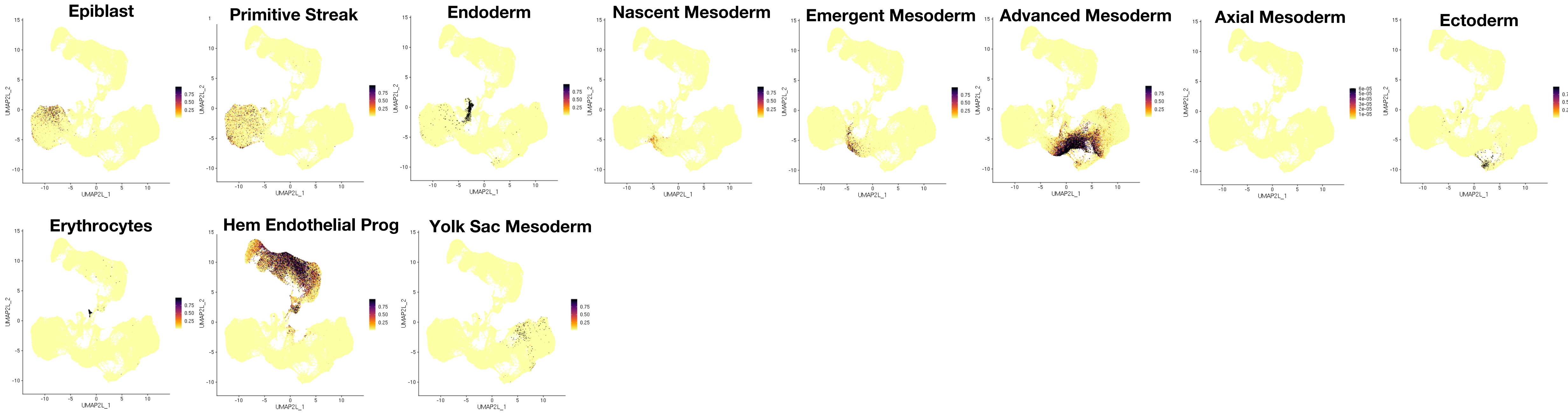


**B** **FETAL LIVER (+kidney +skin) dataset cell types**

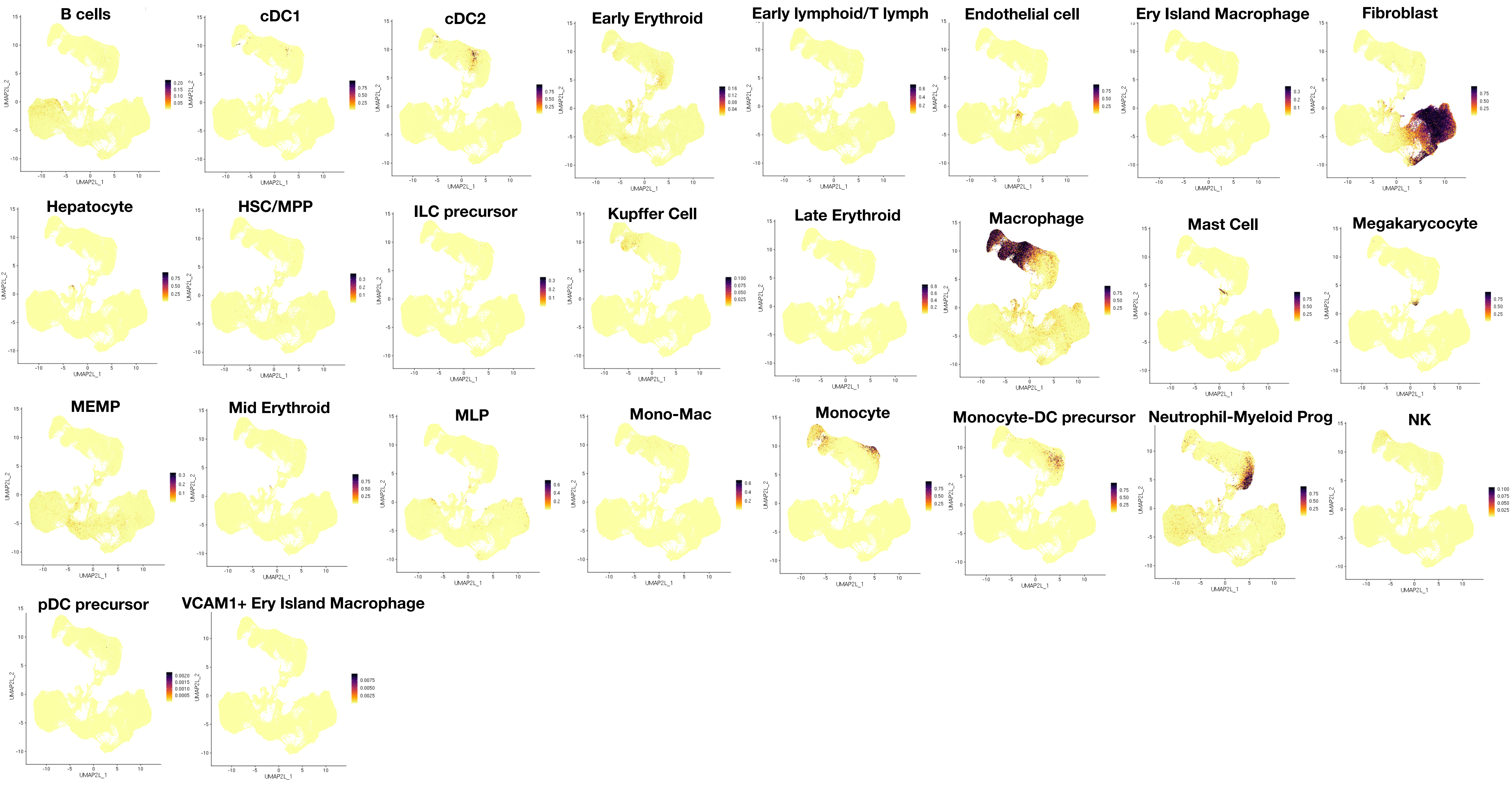


**C** **FETAL YOLK SAC dataset cell types** (not shown in Fig. 1H)

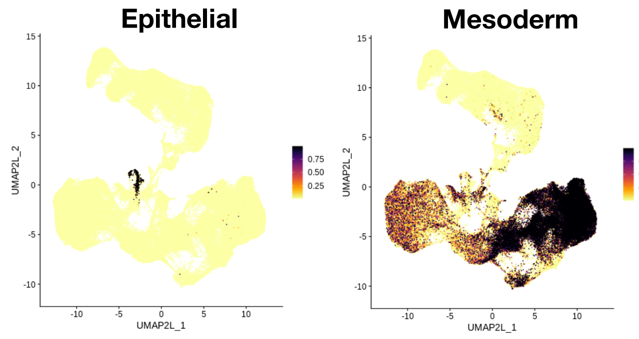

CONTINUED Supplementary Figure 2.

#### D FETAL THYMUS (+ liver) dataset cell types

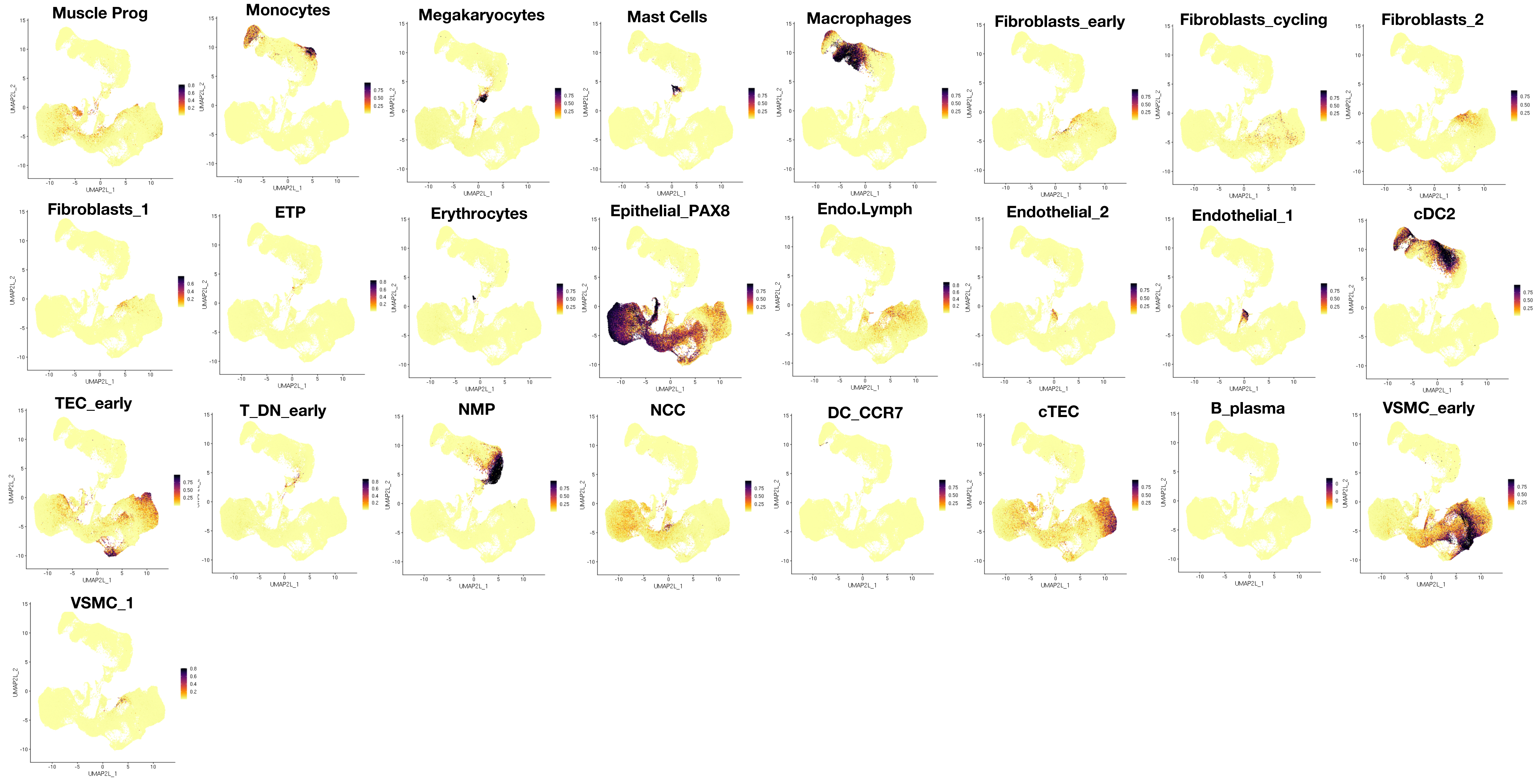


#### E PLACENTA dataset cell types

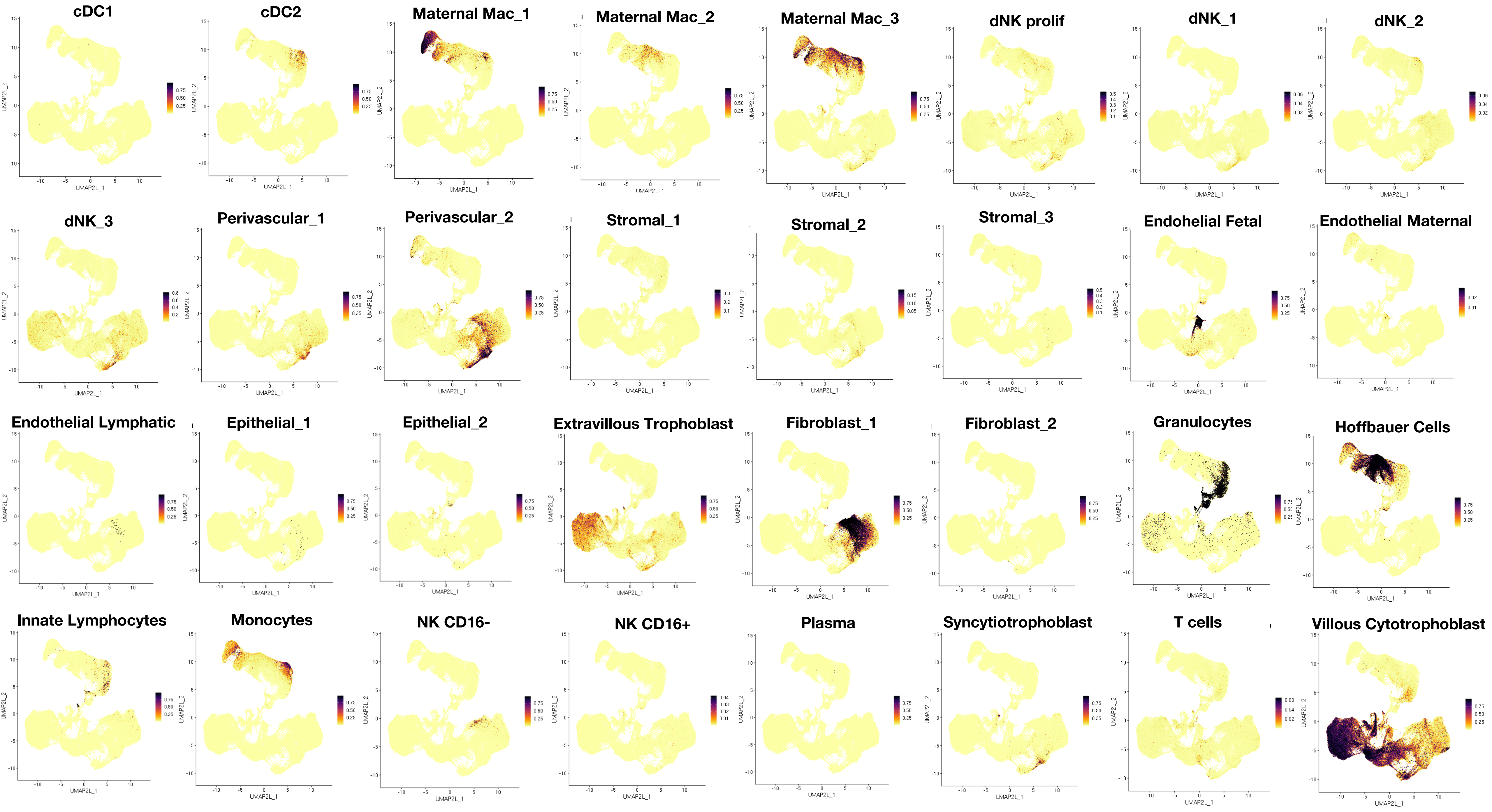

**Supplementary Figure 3.**

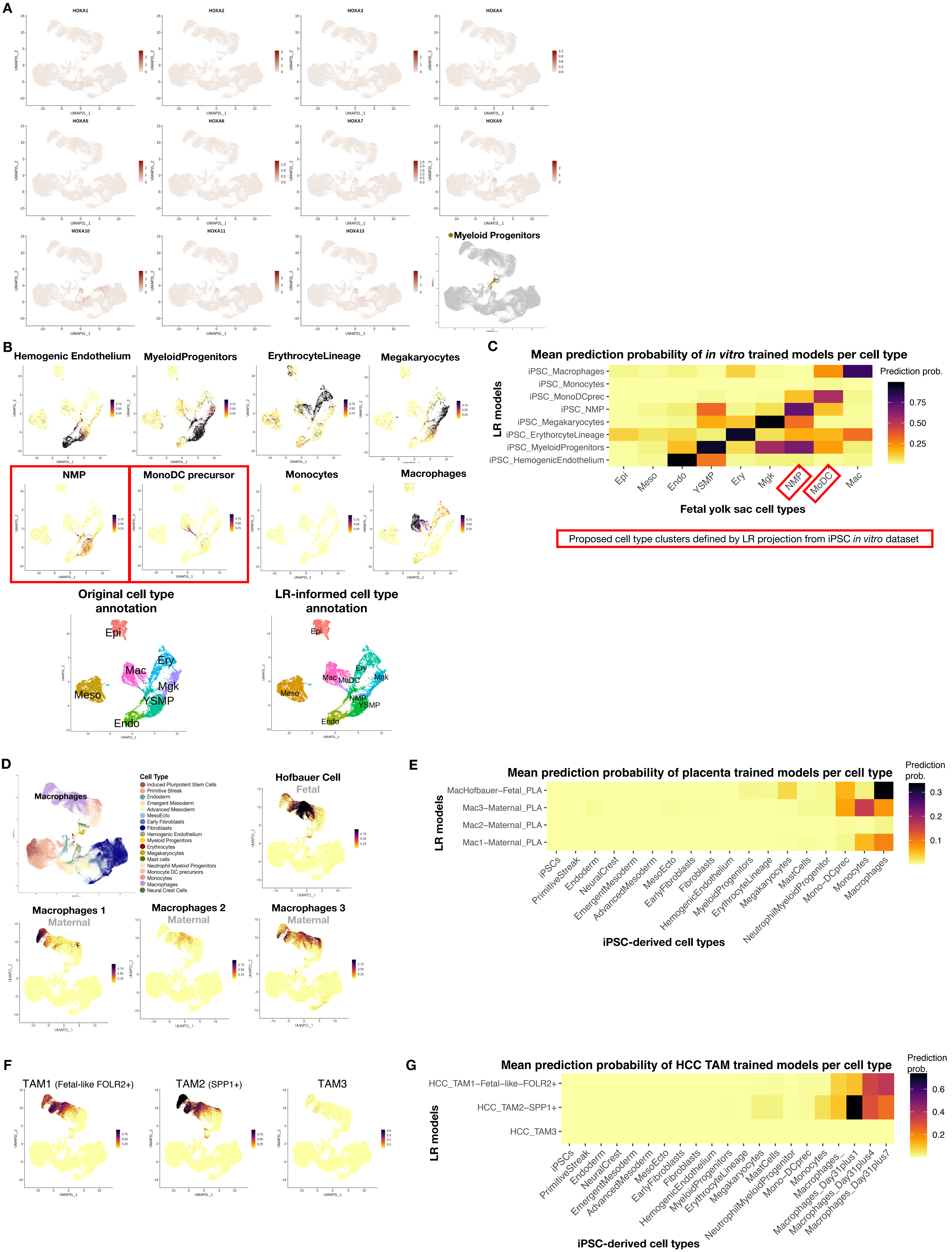

**Supplementary Figure 4.**

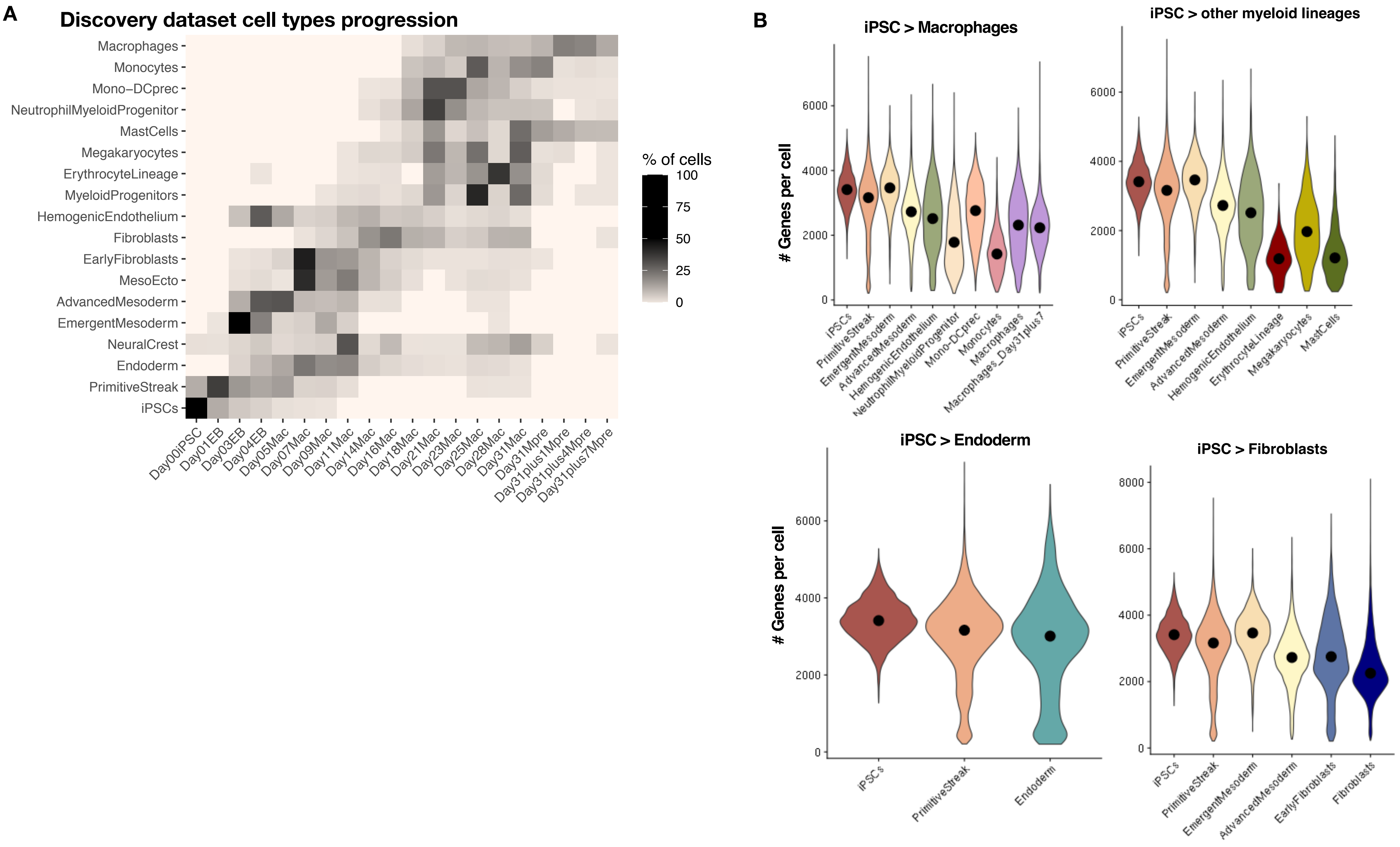

**Supplementary Figure 5.**

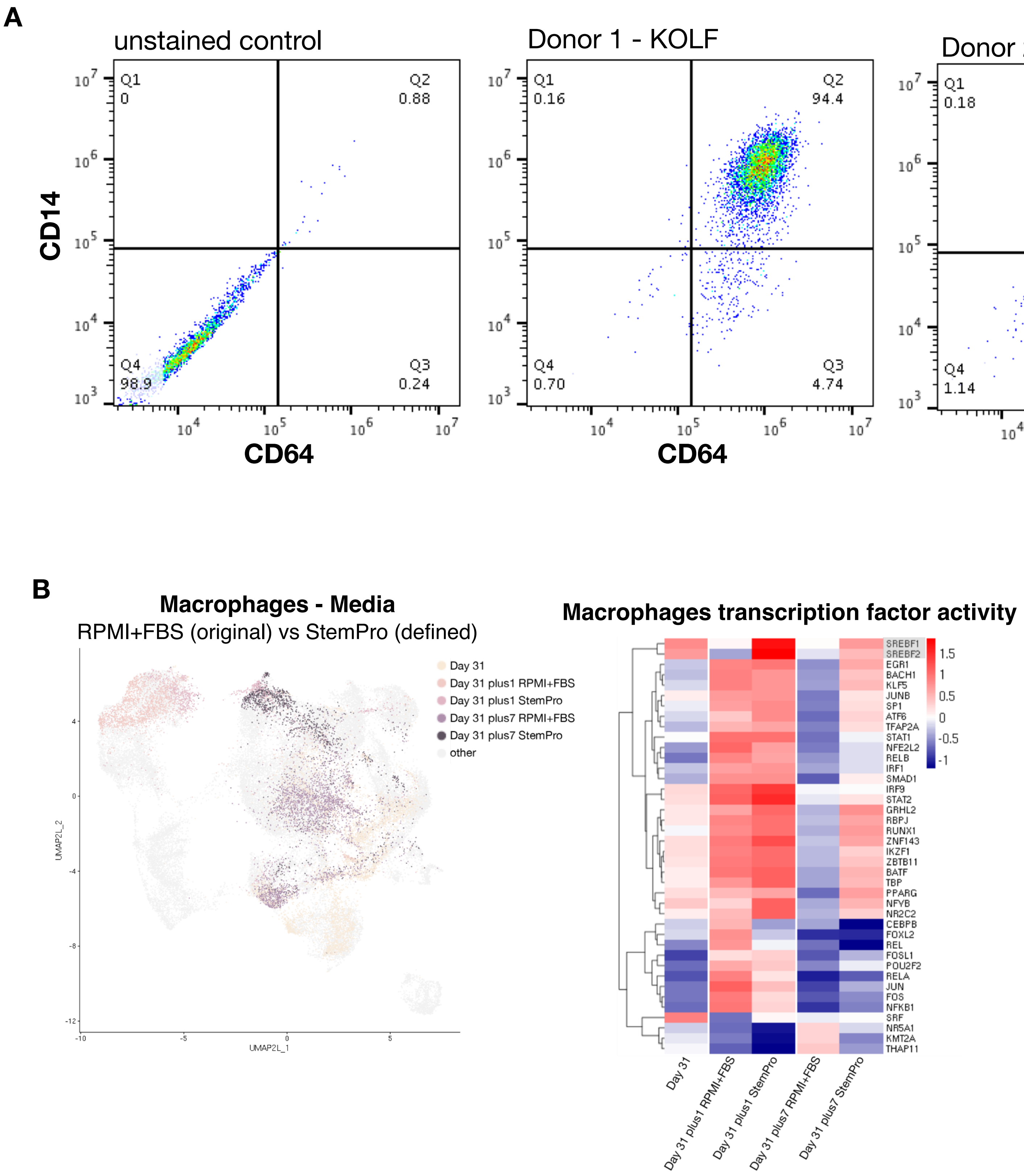

Supplementary Figure 6.

A

#### GASTRULATION dataset

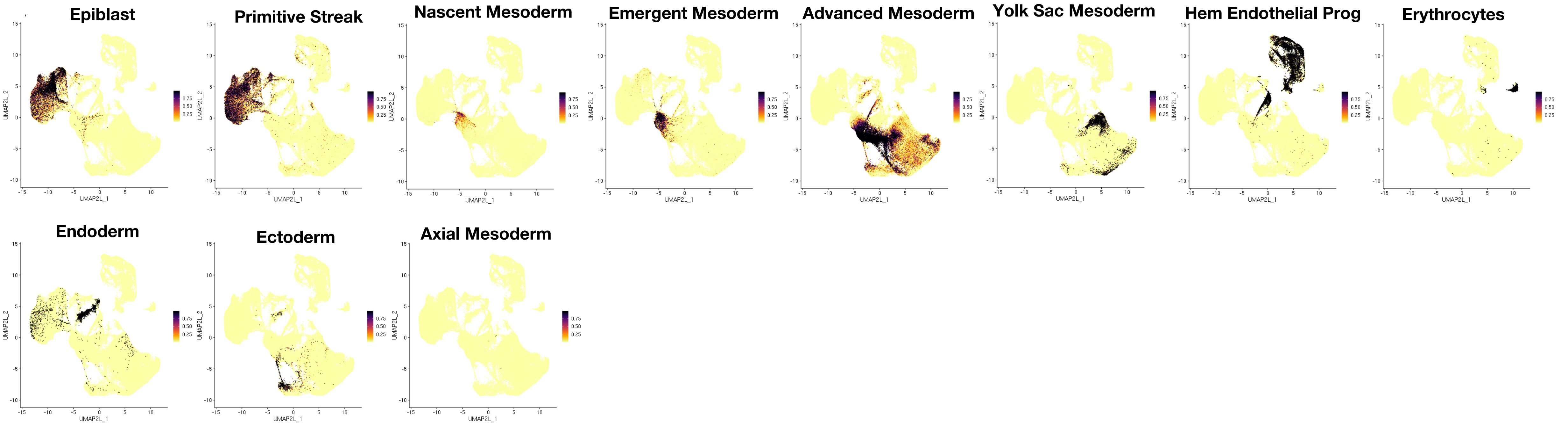


B

#### FETAL LIVER (+kidney +skin) dataset

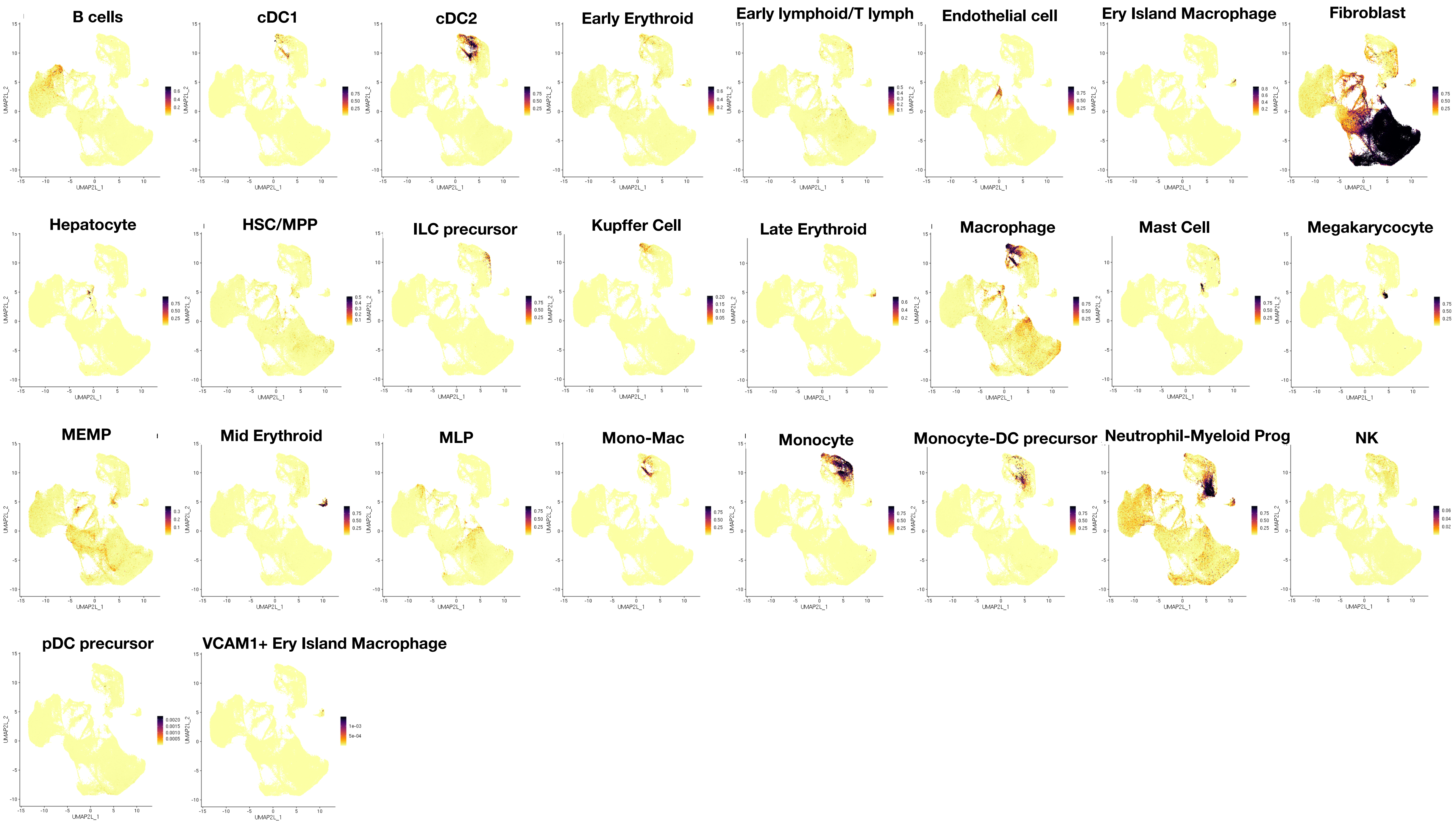


C

#### FETAL YOLK SAC dataset cell types

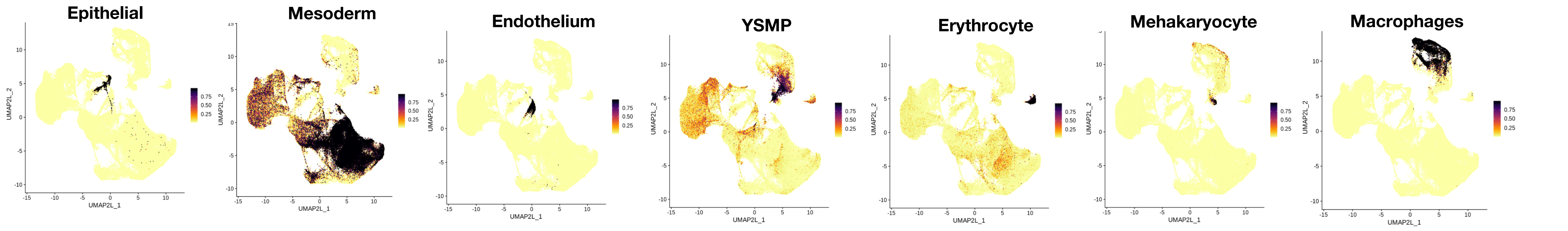

**CONTINUED Supplementary Figure 6.**

**D**

### FETAL THYMUS (+ liver) dataset cell types

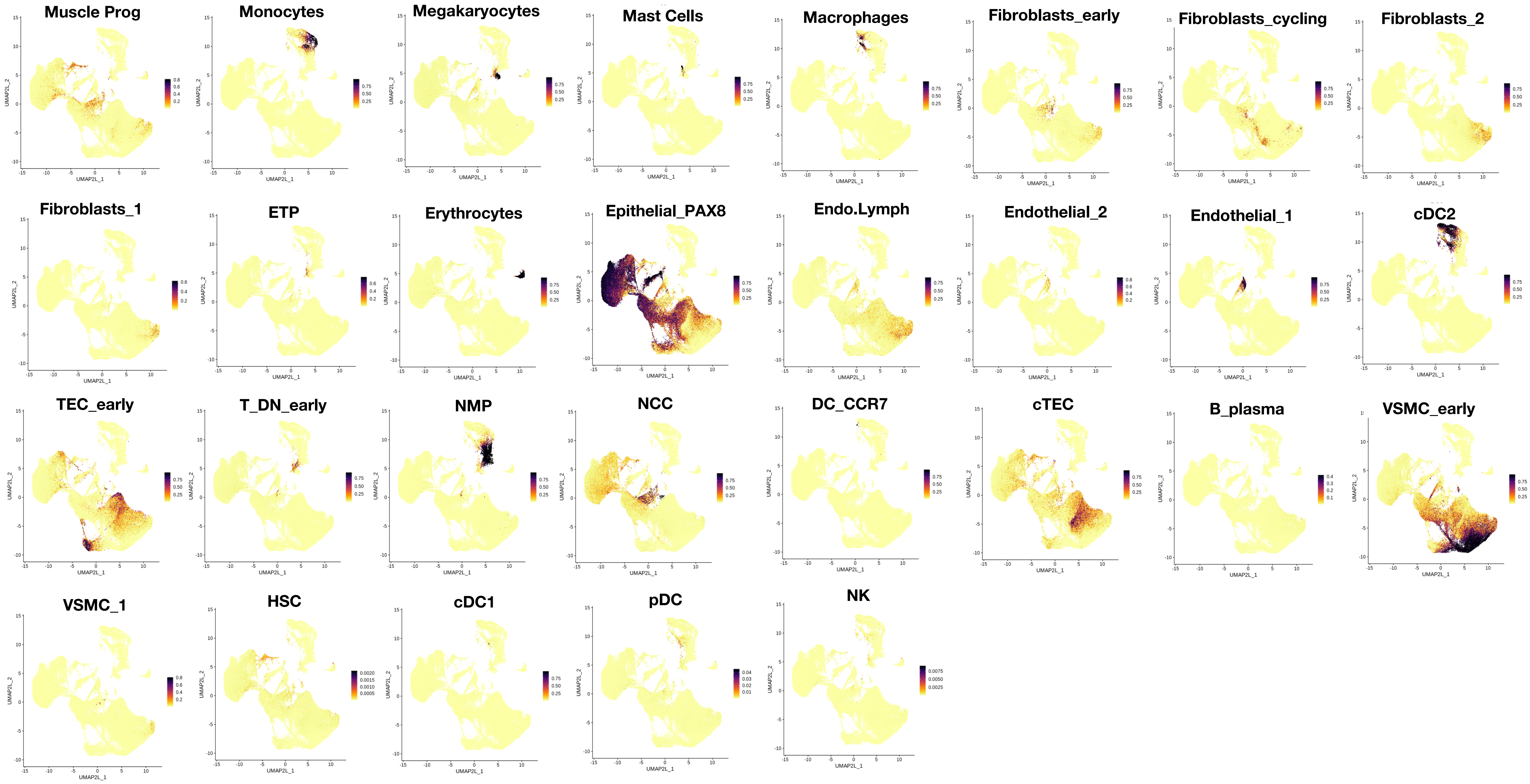

**Supplementary Figure 7.**

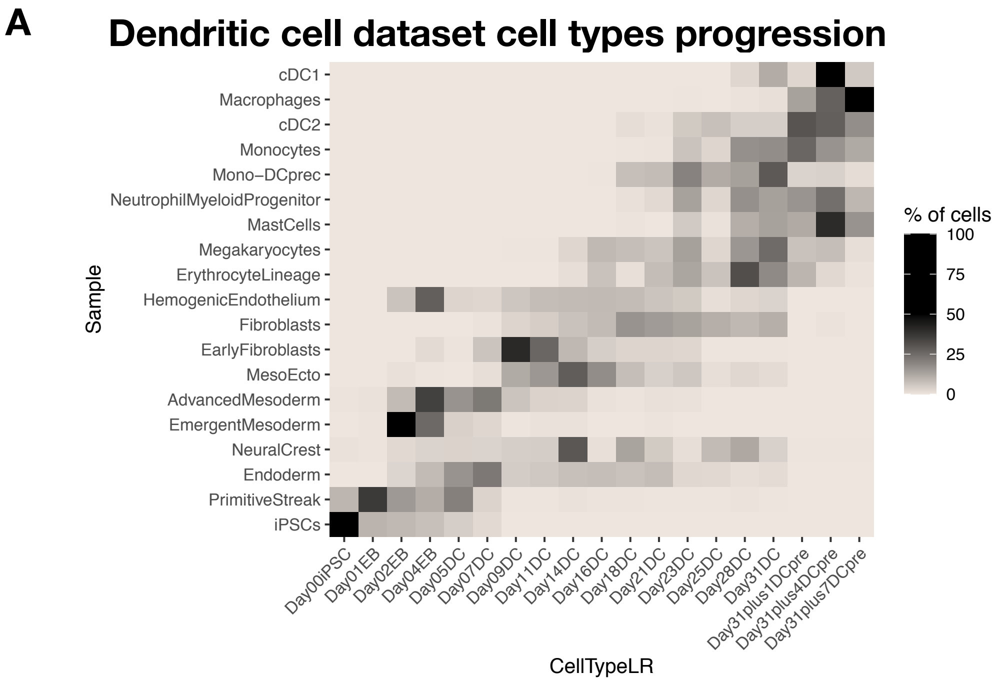

**Supplementary Figure 8.**

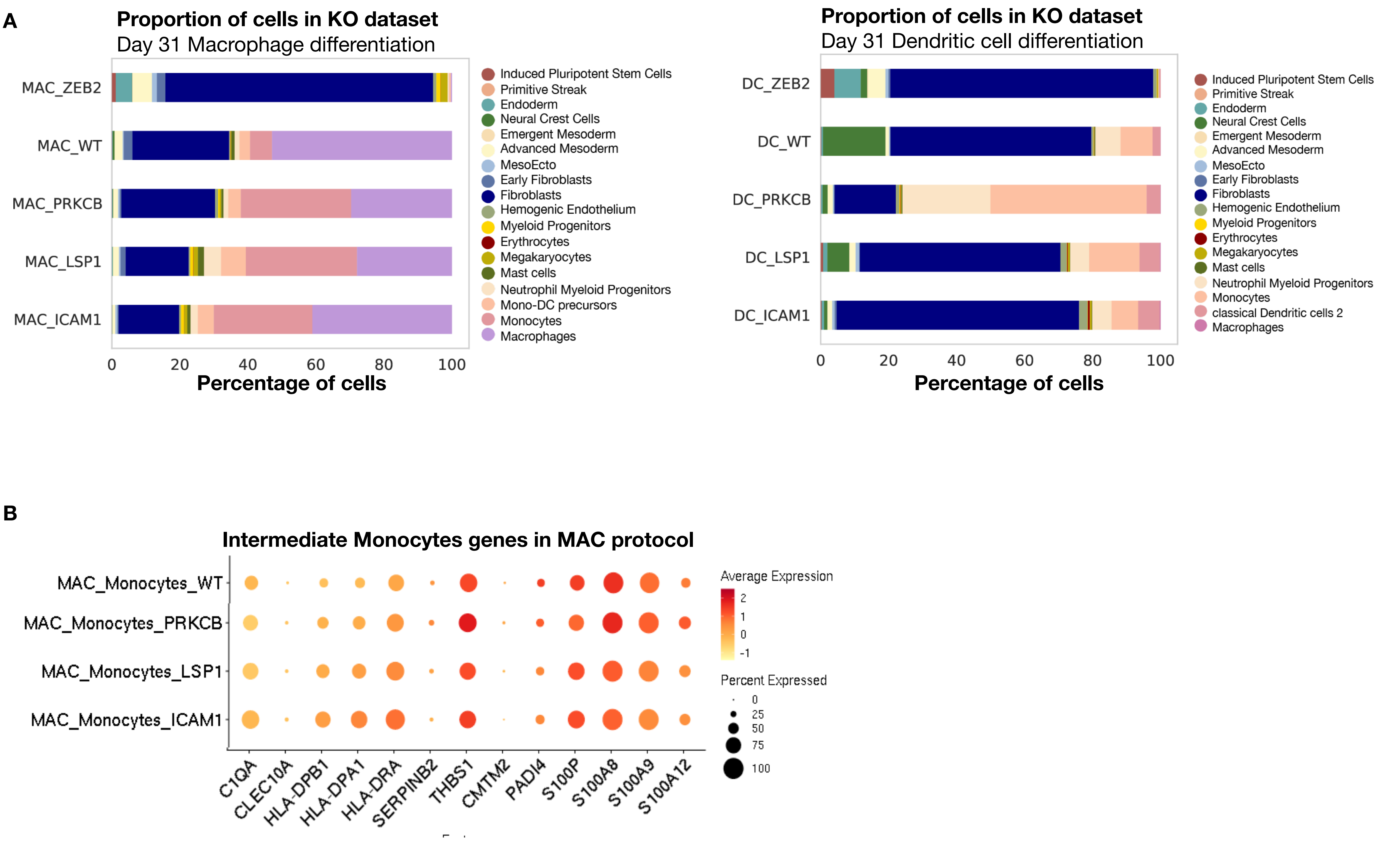
